# A multifaceted approach for obstructive sleep apnea classification from ECG signal using deep learning

**DOI:** 10.1101/2025.09.03.673993

**Authors:** Alan John Varghese, Achilles Gatsonis, Melih Agraz, Vivek Oommen, Anshul Parulkar, Antony Chu, George Em Karniadakis

**Author notes:** these authors contributed equally to this work (Alan John Varghese, Achilles Gatsonis).

## Abstract

Obstructive sleep apnea (OSA) is a common sleep disorder associated with increased cardiovascular and neurocognitive risks. While polysomnography remains the clinical gold standard for diagnosis, it is costly and unsuitable for large-scale or real-time screening. Electrocardiogram (ECG) signals offer a non-invasive, low-cost alternative for sleep apnea detection. We present a *holistic new framework* for OSA detection and forecasting using ECG data on two datasets: PhysioNet Apnea-ECG datasets (healthy patients with apnea), and OSASUD dataset (patients in a stroke unit). Our framework integrates feature engineering methods rooted in dynamical systems theory and statistical analysis. These features are used across a range of models, from conventional machine learning algorithms to novel deep learning architectures. To improve generalization and personalization, we incorporate transfer learning in two ways: across datasets to adapt models trained on large cohorts to smaller clinical datasets, and at the patient level to personalize models using limited individual data, hence demonstrating the use of precision medicine.

## INTRODUCTION

Obstructive sleep apnea (OSA) is a prevalent sleep-related breathing disorder affecting 2-4% of adults globally^1^. OSA is characterized by airway blockage when soft throat tissue collapses during sleep^2^. As oxygen drops and carbon dioxide rises due to the blockage, the patient awakens, sympathetic tone increases, and nasopharyngeal tissue contracts, resulting in a temporary restoration of regular breathing. This cycle often repeats^3^. Cardiovascular effects are particularly notable with the occurrence of sleep disturbances^4,5^, such as apnea. Sustained sympathetic activation from apnea-associated hypoxia can elevated heart rate and blood pressure^6,7^, while inflammatory responses to hypoxia may damage endothelium and promote atherosclerosis^3^. OSA is associated with cardiovascular disease and hypertension^7^, yet it remains underdiagnosed in both men and women. Approximately 85% of patients are undiagnosed, which has marked societal effects due to daytime drowsiness that can lead to missed workdays, vehicle accidents, and workplace accidents^3^. Ultimately, the underdiagnosis of the disease is associated with an annual cost of approximately $3.4 billion^3^. Population-based studies reveal that mild symptoms of sleep apnea affect up to 20% of patients, and up to 7% of patients have moderate to severe sleep apnea^3^. Men are disproportionately affected, with incidence rates two to three times higher than those observed in women^3^. Furthermore, older adults, particularly those between the ages of 68-95, are at a significantly higher risk of developing OSA-related symptoms^3^. Patients who have had a stroke are at an especially high risk of apnea; up to 91.2% of individuals with a stroke have OSA, and 44.6% have a severe manifestation of the disorder^8^.

While at-home tools are emerging^9,10^, diagnosis primarily relies on in-lab polysomnography, which records comprehensive physiological data including EEG (electroencephalogram), EOG (electrooculography), EMG (electromyography), oxygen saturation, and ECG (electrocardiogram)^11^. Polysomnography requires overnight administration by trained technicians and physician interpretation, resulting in high cost^12^. Additionally, the first night and reverse first night effects, as well as instrument cable or electrode malfunction, can result in diagnostic inaccuracies^12–14^. Furthermore, polysomnography is not capable of detecting OSA in real-time, which could be critical in the development of responsive therapies. These limitations emphasize the need for predictive, cost-effective diagnostic tools, particularly for high-risk groups like stroke patients. Electrocardiograms, which have been used to detect arrhythmia and other abnormalities^15,16^, show promise in apnea detection as cardiac activity reflects apneic events. Heart rate variability spectrograms show cardiopulmonary coupling mode shifts between apneic and non-apneic states^17^. Healthy patients primarily exhibit high frequency coupling, whereas unhealthy patients exhibit low frequency coupling^17^. Sleep apnea also results in sympathetic overdrive, producing heart rate variability changes that can be detected on an electrocardiogram^18^. These relationships between cardiac activity and apnea have made electrocardiogram-based data valuable for machine learning-based apnea detection. Techniques including boosting methods^19,20^, support vector machines^21,22^, discriminant analyses^23^, k-nearest neighbors^24^, convolutional neural networks^25^, gated recurrent unit networks^25,26^, long short-term memory models^27,28^, and transformer models^29,30^ have been used to detect apnea using ECG data with accuracies ranging from over 80 to over 90%, validating the applicability of ECG in apnea detection.

The latest models utilize transformer-based architectures, originally designed for natural language processing, to detect apnea based on time series ECG data^29,30^ to obtain detection accuracies over 90%. However, it is unclear why such transformer-based architectures perform well, and performance varies across datasets^30^. Furthermore, studies most often are performed on the PhysioNet Apnea-ECG Database^11^, which consists of heavily-selected nighttime ECG recordings from healthy patients, leaving gaps in research on populations with comorbidities like stroke. Different datasets also vary in apnea/non-apnea label ratios, resulting in possible data imbalance issues. Additionally, equipment variability between hospitals may introduce inconsistent signal quality. The imbalance issue has been previously addressed in classification problems using data augmentation methods, including mixup^31^, synthetic minority oversampling technique (SMOTE)^32^, and Gaussian noise. Mixup and SMOTE are resampling methods that address the issue of data imbalance by providing more samples of the minority class, whereas Gaussian noise alleviates the issue of overfitting to training data. Furthermore, while Bahrami and Forouzanfar^25^ used recurrent neural networks (RNN) to forecast apneic activity, most studies have focused on detection of past apneic events rather than forecasting future apneas. This limits models to diagnostic settings, whereas forecasting models could be used to therapeutically prevent apnea. Finally, machine learning approaches often select ECG-derived features that detect apnea but could lack clinical interpretability, resulting in black-box models.

In this study, we use a multifaceted approach integrating artificial intelligence, statistical analyses, and system dynamics to detect and forecast apnea using ECG data, demonstrated in Fig. 1. We analyze system dynamics through the kurtosis, which provides information about the intermittency of the ECG time series, the Lyapunov exponent, which measures how chaotic the system is, and wavelet-based analyses, which decomposes ECG features using continuous wavelet transforms, in conjunction with the occurrence of apnea. Statistical analyses, namely correlational studies of ECG-derived features and the presence of disease, are used to assess the clinical significance of ECG-derived features. Additionally, artificial intelligence-based feature selection methods, such as Shapley Additive exPlanations (SHAP) feature importance^33^ is used to assess the importance of features in the determination of apnea versus non-apnea. We leverage recent advances in machine learning, particularly transformer-based architectures, as well as more conventional approaches including logistic regression, k-nearest neighbors, extreme gradient boosting, LSTM, and convolutional neural networks, to detect apnea and compare the viability of different algorithms in the detection of apnea. We also use a predictive model to forecast future occurrence of sleep apnea. We apply our approach both on healthy patients with no comorbidities (PhysioNet Apnea-ECG Database^11^), as well as for patients admitted to a hospital for a stroke (Obstructive Sleep Apnea Stroke Unit Dataset (OSASUD)^8^). We provide a method for electrocardiogram data preprocessing and feature engineering applicable across different datasets. We also present an integration of statistical significance and artificial-intelligence feature importance to select feature sets that are of both clinical significance and model significance, supporting high performance and high clinical interpretability. Our assessment of diverse patient populations, specifically healthy patients^11^ and patients with stroke^8^ represents a new approach in apnea detection from electrocardiograms and broadens the scope of the field to patients of different health backgrounds. Our forecasting of apneic activity offers a novel contribution to the field, which largely emphasizes detection. Lastly, we apply transfer learning methods to the St. Vincent’s University Hospital / University College Dublin Sleep Apnea Database^34^, demonstrating applicability of our model to unseen data, ensuring robustness across patient populations. Our combined methodology addresses apnea detection and prediction across patient groups and aims for reproducibility, justifiable processing, and clinically meaningful model development.

**Figure 1.**
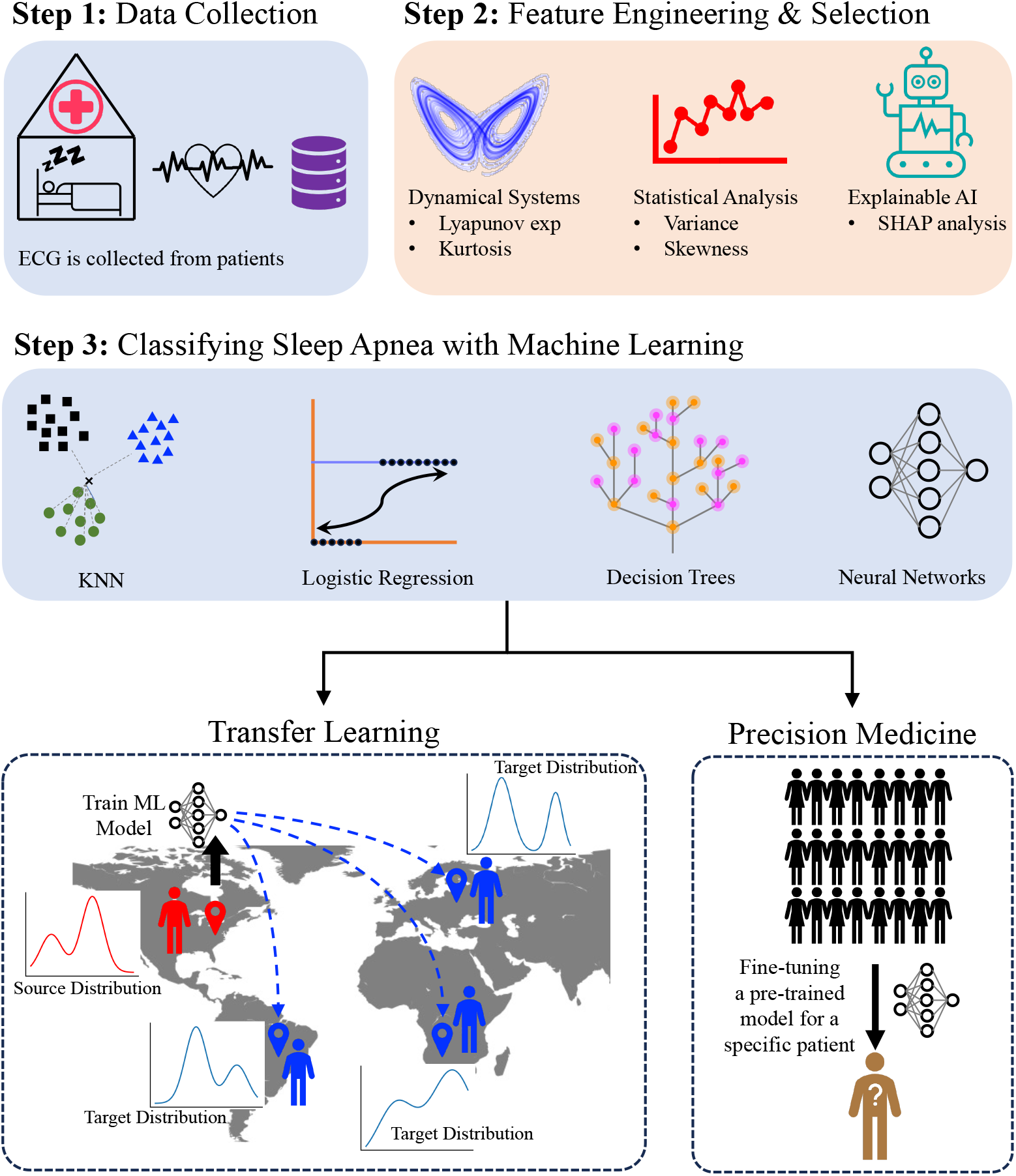
An illustrative overview of the proposed framework. We utilize publicly available ECG datasets with annotated apnea events. Feature engineering, inspired by dynamical systems and statistical analysis, is applied to derive relevant features from the ECG signals. Both classical machine learning (ML) and deep learning models are employed to classify apnea events, with the best-performing deep learning model selected for further tasks, such as forecasting and diagnosing the severity of obstructive sleep apnea (OSA). Additionally, we explore transfer learning strategies to enhance model performance in datasets with limited data as well as improve personalized predictions.

## RESULTS

### Feature Engineering

In this study, two primary datasets are employed: PhysioNet Apnea-ECG and OSASUD. After preprocessing, the PhysioNet Apnea-ECG dataset includes 16,810 observations with 80 variables, while OSASUD has 7,609 observations and 85 variables, with 78 features shared between them. PhysioNet includes basic demographic data (e.g., *Height, Weight*), whereas OSASUD incorporates advanced physiological markers such as BMI, HRV metrics (HFD, KFD, MSEn), Lyapunov Exponent, and 10-Minute Kurtosis. To enhance the data’s representational power, we extracted both statistical and time-series features from ECG signals. This included standard metrics (`mean, skewness, lagged values`) and rolling window transformations (`ECG_Mean_Rolling, ECG_Skew_Rolling`) computed from one-minute ECG segments, particularly in the OSASUD dataset. A detailed list of features used for the two datasets is provided in Tables 4 and 5 in Supplementary Materials. This feature engineering pipeline provides a robust statistical representation of ECG dynamics, supporting the identification of key OSA indicators. These features were incorporated into machine learning models to improve apnea prediction and model interpretability.

Figure 2 (a, b) presents the SHAP values for the 15 most influential features from the PhysioNet Apnea-ECG and OSASUD datasets. These analyses illustrate which features the classification models rely on most when distinguishing between apnea and non-apnea conditions. In the PhysioNet dataset, the most significant variables are primarily statistical and temporal features derived from the ECG signal (e.g., ECG Mean, RR Skw, HRV MSEn). In contrast, the OSASUD dataset highlights features such as the Lyapunov exponent, HRV entropy measures (MSEn 5, HFD), and higher-order ECG features (ECG Kurtosis). This suggests that the OSASUD dataset exhibits more complex physiological characteristics and more pronounced effects of comorbidities. A particularly noteworthy finding is that statistical moments of the ECG signal (such as kurtosis, skewness, and mean) exhibit high SHAP values in both datasets. This indicates that structural alterations in the cardiac signal, associated with apnea, are prominent and can be effectively modeled.

**Figure 2.**
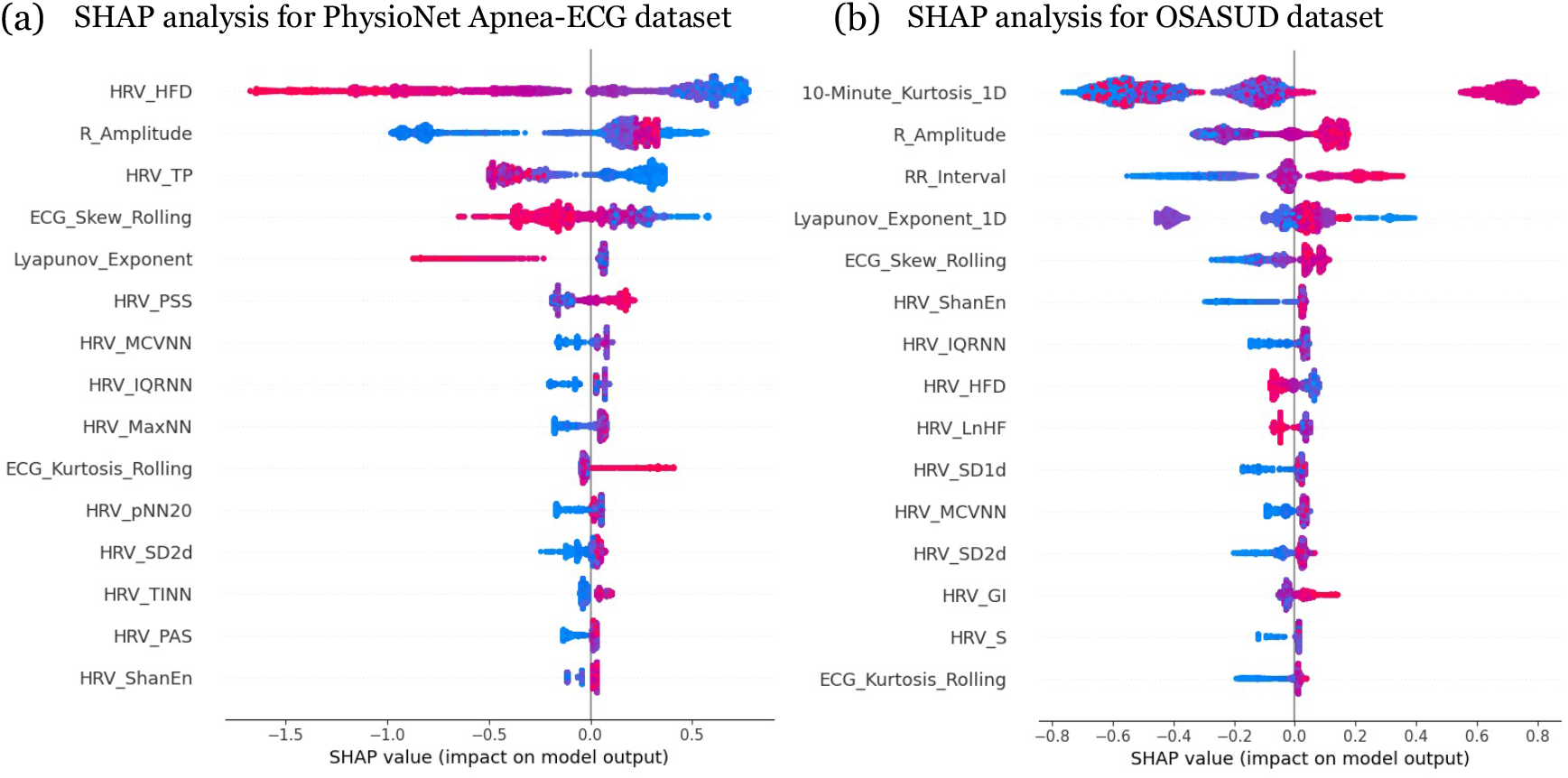
SHAP Analysis of Feature Importance. The plots illustrate the 15 most influential features, as determined by SHAP values, for models classifying apnea and non-apnea events in the (a) PhysioNet Apnea-ECG and (b) OSASUD datasets. (a) For the PhysioNet dataset, the analysis highlights the strong influence of features like HRV_HFD, R_Amplitude, and HRV_TP andLyapunov_Exponent. Other contributing features include various HRV metrics (e.g., HRV_PSS, HRV_MCVNN, HRV_IQRNN) and statistical measures of the ECG (e.g., ECG_Skew_Rolling, ECG_Kurtosis_Rolling) (b) In the OSASUD dataset, 10-Minute_Kurtosis_1D, R_Amplitude, RR_Interval, and Lyapunov_Exponent_1D are identified as key predictors. The feature importance profile also includes several HRV measures (e.g., HRV_ShanEn, HRV_IQRNN, HRV_HFD) and statistical ECG features (e.g., ECG_Skew_Rolling, ECG_Kurtosis_Rolling), indicating a more distributed influence across feature types compared to the PhysioNet dataset.

These features, which influence the model’s decision-making process, also carry clinical significance. Chattopadhyay et al. (2018)^35^ utilized kurtosis analysis based on Discrete Wavelet Transform (DWT) for diagnosing sleep apnea, reporting significant differences in kurtosis values derived from ECG signals of apneic patients versus healthy individuals. Smbatovna and Aleeksevna^36^ demonstrated that although the Lyapunov exponent is effective in evaluating chaotic dynamics in EEG signals, it yields an accuracy of approximately 90%, underscoring its value as a supportive analytical tool. Variations in R-wave amplitude have been explored as a potential method for identifying sleep apnea cases^37^. HRV analysis has also been proposed as a screening approach for obstructive sleep apnea (OSA)^38^. Moreover, studies have shown that, in comparison to healthy individuals, reduced heart rate complexity—assessed through entropy-based measures—can serve as a sensitive marker for the detection of OSA^39,39,40^.

### Dynamical Systems Perspective

We compute the largest Lyapunov exponent and kurtosis of the ECG signal for patients in both the PhysioNet Apnea-ECG dataset and the OSASUD dataset. The largest Lyapunov exponent can be used to assess the chaotic nature of a dynamical system. The kurtosis quantifies the signal’s intermittency by measuring its ‘tailedness’, thereby indicating how often outliers occur. In Fig. 3 (a, c), we present the variation of the largest Lyapunov exponent over time alongside the occurrence of apnea events (shaded in blue) for patients in both datasets. The Lyapunov exponent tends to increase during periods of non-apnea and decrease during apnea, consistent with previous studies: Han and Wang^41^ found that the average largest Lyapunov exponent of patients with arrhythmia are lower than that of healthy individuals, and Tayel and AlSaba^42^ demonstrated that patients with congestive heart failure exhibit lower Lyapunov exponents compared to healthy controls. Thus, our results support the notion that a reduction in the largest Lyapunov exponent of the ECG signal is associated with disease states. In Fig. 3 (b, d) we present the kurtosis and the apnea occurrence profiles for patients in both datasets. The kurtosis tends to decrease during periods of non-apnea and increases during apnea, indicating greater intermittency during apnea. This trend contrasts with the behavior of the Lyapunov exponent, which decreases during apnea.

**Figure 3.**
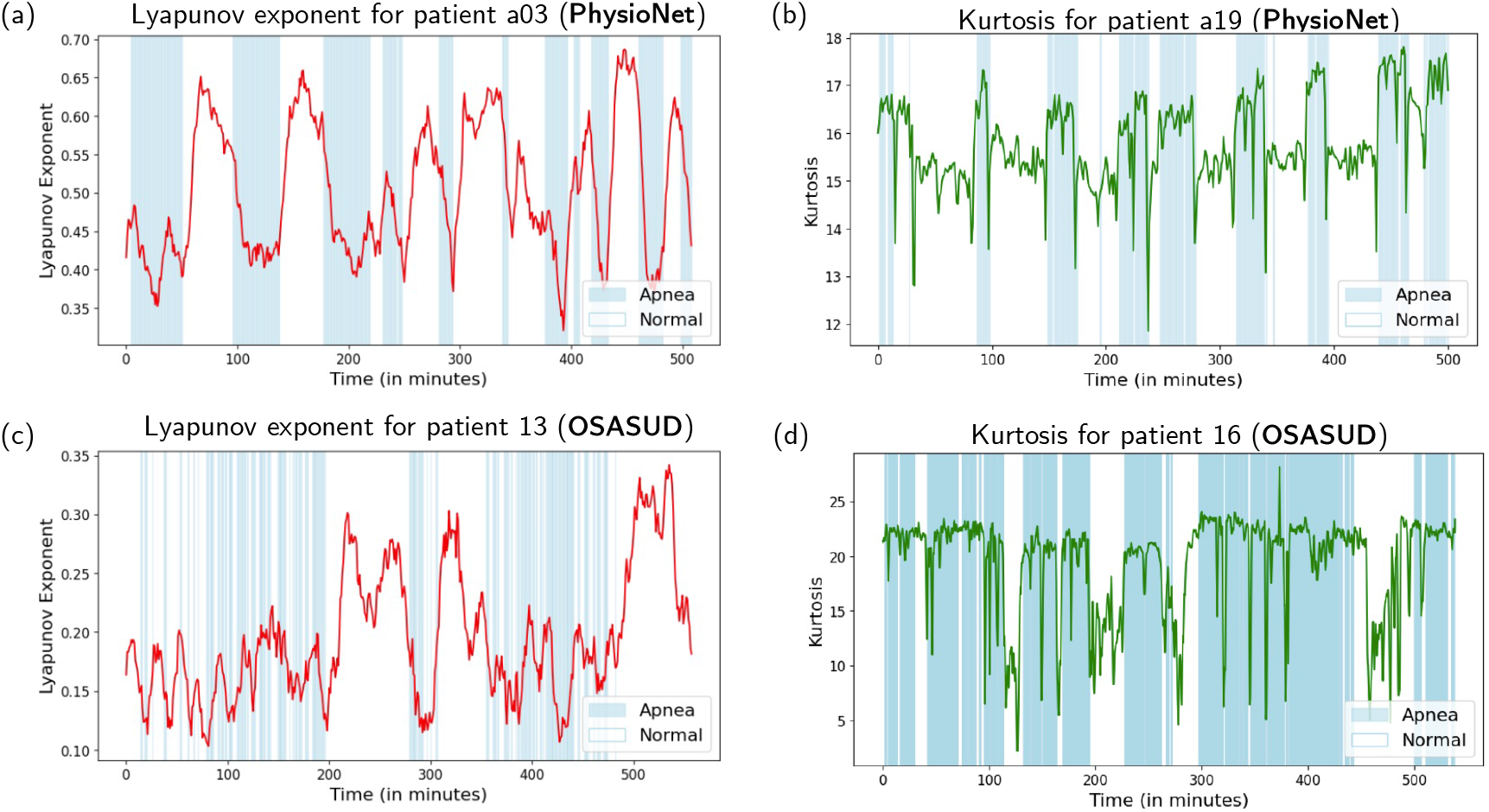
Correlation of Lyapunov exponent and kurtosis with patient conditions in PhysioNet Apnea-ECG and OSASUD datasets. We show the variation of (**a**,**c**) Lyapunov exponent and (**b**,**d**) kurtosis with time for two representative patients in PhysioNet Apnea-ECG dataset and OSASUD dataset, respectively. The Lyapunov exponent tends to decrease during apnea states (blue), and increases during normal states (white). In contrast, in the case of kurtosis we see the opposite trend. For clarity of representation, we have chosen the patients with the highest anti-correlation (−0.72 and −0.21) in the case of Lyapunov exponent (patient a03 and patient 13), and the patients with highest correlation (0.62 and 0.30) in the case of kurtosis (patient a19 and patient 16).

We compute the average correlation coefficients between apnea occurrence and the assessed features for patients in both datasets. In the PhysioNet Apnea-ECG dataset, we observe a stronger negative correlation (−0.20) between the Lyapunov exponent and apnea compared to the OSASUD dataset (−0.02). Similarly, we observe a stronger positive correlation (0.16) between kurtosis and apnea in the PhysioNet Apnea-ECG dataset compared to the OSASUD dataset (0.07). The lower correlations in the OSASUD dataset are likely due to the increased complexity of this dataset, which includes the presence of other comorbidities in the patients.

### Model Performance

#### Conventional ML Models

Initially, the models were assessed based on **Three Features**—R amplitude, RR interval, and the first-order difference of the RR interval—forming a minimal feature set inspired by Hu et al.^29^. Next, they were evaluated using **All Features** in the dataset, including static covariates. Following this, the models were tested with **Significant Features** that showed statistical differences between patients with and without apnea. Ultimately, the final feature set, referred to as **Significant ECG Features**, consisted of only the statistically significant features, excluding static covariates, resulting in a feature set derived solely from ECG data. All conventional results were computed across these four feature sets, providing a thorough assessment of model performance under varying feature configurations, as presented in Table 1.

**Table 1.** Performance metrics of conventional machine learning models to detect sleep apnea in the PhysioNet Apnea-ECG and OSASUD dataset.

| Dataset | Approach | Feature Set | Sens. | Spec. | Acc. | Prec. | F1 score |
| --- | --- | --- | --- | --- | --- | --- | --- |
| PhysioNet | LR | Three Features | 0.013 | **0.993** | 0.619 | 0.538 | 0.030 |
|  | LR | All Features | 0.678 | 0.817 | 0.764 | 0.695 | 0.690 |
|  | LR | Significant Features | 0.639 | 0.826 | 0.755 | 0.693 | 0.670 |
|  | LR | Significant ECG Features | 0.444 | 0.824 | 0.679 | 0.608 | 0.510 |
|  | XGBoost | Three Features | 0.273 | 0.758 | 0.573 | 0.411 | 0.330 |
|  | XGBoost | All Features | 0.636 | 0.877 | 0.786 | 0.762 | 0.690 |
|  | XGBoost | Significant Features | 0.656 | 0.890 | 0.801 | 0.786 | 0.710 |
|  | XGBoost | Significant ECG Features | **0.707** | 0.896 | **0.807** | 0.769 | **0.740** |
|  | RF | Three Features | 0.303 | 0.755 | 0.583 | 0.433 | 0.360 |
|  | RF | All Features | 0.639 | 0.904 | 0.803 | **0.804** | 0.710 |
|  | RF | Significant Features | 0.610 | 0.898 | 0.788 | 0.788 | 0.690 |
|  | RF | Significant ECG Features | 0.647 | 0.896 | 0.801 | 0.793 | 0.710 |
| OSASUD | LR | Three Features | 0.000 | **0.999** | **0.835** | 0.000 | 0.000 |
|  | LR | All Features | 0.004 | 0.995 | 0.833 | 0.143 | 0.008 |
|  | LR | Significant Features | 0.002 | 0.995 | 0.831 | 0.053 | 0.003 |
|  | LR | Significant ECG Features | 0.000 | **0.999** | **0.835** | 0.000 | 0.000 |
|  | XGBoost | Three Features | **0.088** | 0.948 | 0.807 | 0.249 | **0.130** |
|  | XGBoost | All Features | 0.011 | 0.974 | 0.816 | 0.075 | 0.019 |
|  | XGBoost | Significant Features | 0.016 | 0.978 | 0.820 | 0.128 | 0.029 |
|  | XGBoost | Significant ECG Features | 0.049 | 0.960 | 0.811 | 0.196 | 0.079 |
|  | RF | Three Features | 0.029 | 0.988 | 0.831 | **0.325** | 0.054 |
|  | RF | All Features | 0.001 | **0.999** | **0.835** | 0.167 | 0.002 |
|  | RF | Significant Features | 0.001 | 0.998 | 0.834 | 0.062 | 0.002 |
|  | RF | Significant ECG Features | 0.005 | 0.997 | 0.834 | 0.269 | 0.011 |

#### Deep Learning Models

We evaluated the performance of two deep learning models - Hybrid Transformer and ApneaFormer - to detect apnea instances, aiming to address limitations observed with conventional ML methods, particularly on the challenging OSASUD dataset. As outlined in the Methods section, both the models were trained on three-minute segments of ECG data to predict the presence of apnea in the central minute. In addition to the ECG signal, we explore augmenting the input features with RR interval, kurtosis and Lyapunov exponent to improve performance, particularly on the more challenging OSASUD dataset.

Table 2 summarizes the results. On the PhysioNet Apnea-ECG dataset, ApneaFormer consistently outperformed all baseline methods, achieving an F1 score of 0.914 and AUC of 0.981 using ECG alone. Notably, adding extra features did not yield noticeable gains on this dataset, possibly due to a performance plateau already achieved using ECG alone (>90% in all evaluation metrics). The training and testing splits followed the standard protocol of using the first 35 patients for training and the remaining 35 for testing.

**Table 2.**
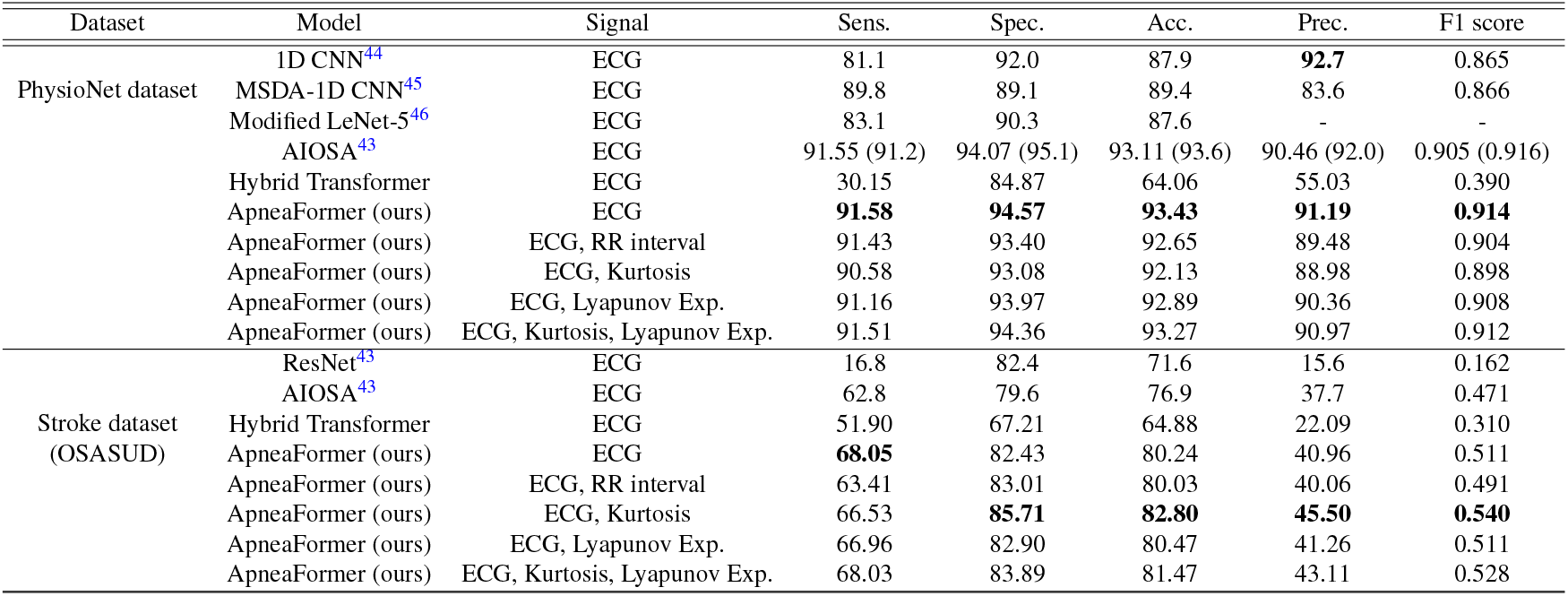
Performance metrics of our ApneaFormer model on the PhysioNet-Apnea ECG dataset and OSASUD dataset. We evaluate the performance of our model using different feature combinations and compare it against other baseline models.

In contrast, the OSASUD dataset presented additional challenges, including the presence of comorbidities and increased signal noise. Convention ML models performed poorly for this dataset. ApneaFormer helped to improve the performance substantially. Including additional features such as kurtosis and Lyapunov exponent led to further gains. We obtained peak performance when using ECG signal along with kurtosis, achieving an F1 score of 0.540. These results were obtained using leave-one-out cross-validation across all 30 patients, consistent with the evaluation strategy in Bernardini et al.^43^.

Across both datasets, ApneaFormer consistently outperformed the Hybrid Transformer. We attribute this to ApneaFormer’s ability to process the ECG signal at its original resolution, while the Hybrid Transformer applies aggressive downsampling, likely leading to a loss of relevant temporal information. Details about the model architecture, training setup and hyperparameters are provided in the Methods section. Although, we attempted to reproduce the AIOSA results using the author’s official implementation, our experiments attained slightly lower performance than originally reported (values in the parantheses). We include these results for transparency and consistency in benchmarking.

Overall, our results demonstrate that deep learning methods, particularly ApneaFormer, substantially improve performance on complex, real-world datasets such as OSASUD, while maintaining state-of-the-art performance on benchmark datasets like PhysioNet Apnea-ECG.

##### ROC Curve

Figure 4 (a, b) shows the Receiver Operating Characteristic (ROC) curves for our ApneaFormer model evaluated on the PhysioNet Apnea-ECG dataset and the OSASUD dataset. The ROC curve plots the true positive rate (sensitivity) against the false positive rate across varying classification thresholds. A model with better performance yields a curve that rises steeply towards the top-left corner, indicating higher sensitivity with fewer false positives. The area under the curve (AUC) quantifies this: a perfect classifier achieves an AUC of 1.0, while a random classifier has an AUC of 0.5 (indicated by the dashed gray line). Our model achieves a high AUC of 0.981 on the PhysioNet Apnea-ECG dataset and 0.806 on the OSASUD dataset, significantly outperforming a random classifier on both tasks. The lower AUC on the OSASUD dataset reflects the increased difficulty of this dataset, which contains patients with several comorbidities and noisier signals - factors that make classification of apnea from ECG signal challenging.

**Figure 4.**
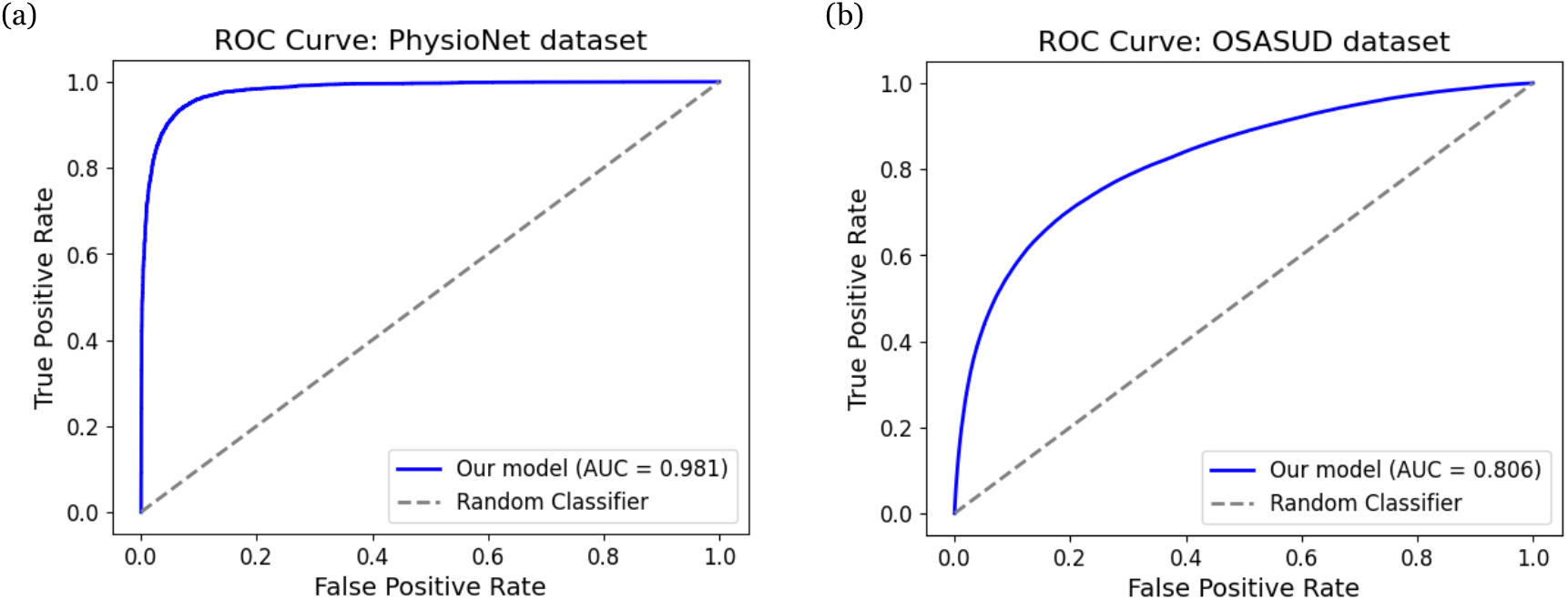
Receiver Operating Characteristic (ROC) curve for the ApneaFormer model on (a) PhysioNet Apnea-ECG dataset and (b) OSASUD dataset. The ROC curve for the PhysioNet Apnea-ECG dataset has a higher area under the curve (AUC = 0.981) compared to the OSASUD dataset (AUC = 0.806), reflecting the increased difficulty and complexity of the OSASUD dataset.

##### Forecast Performance

Figure 5 (a, b) presents the forecast performance of our model on the PhysioNet Apnea-ECG and OSASUD datasets. The model is trained using 3-minute segments of ECG signals as input to predict the occurrence of apnea events at various future time points. We report five evaluation metrics - sensitivity, specificity, accuracy, precision and F1 score - on the test set across forecast horizons from 1 to 30 minutes. For both datasets, there is a consistent decline in performance as the forecast horizon increases. This degradation reflects the inherent challenge of long range forecasting. Comparing the two datasets, we observe that the model performs consistently better on the PhysioNet Apnea-ECG dataset across all metrics and time horizons. This performance gap can be attributed to the greater complexity of OSASUD dataset, which includes patients with multiple comorbities and noisier ECG signals. These results highlight two important aspects: i) the limitations of forecasting further into the future, and ii) the impact of dataset complexity on the models predictive performance.

**Figure 5.**
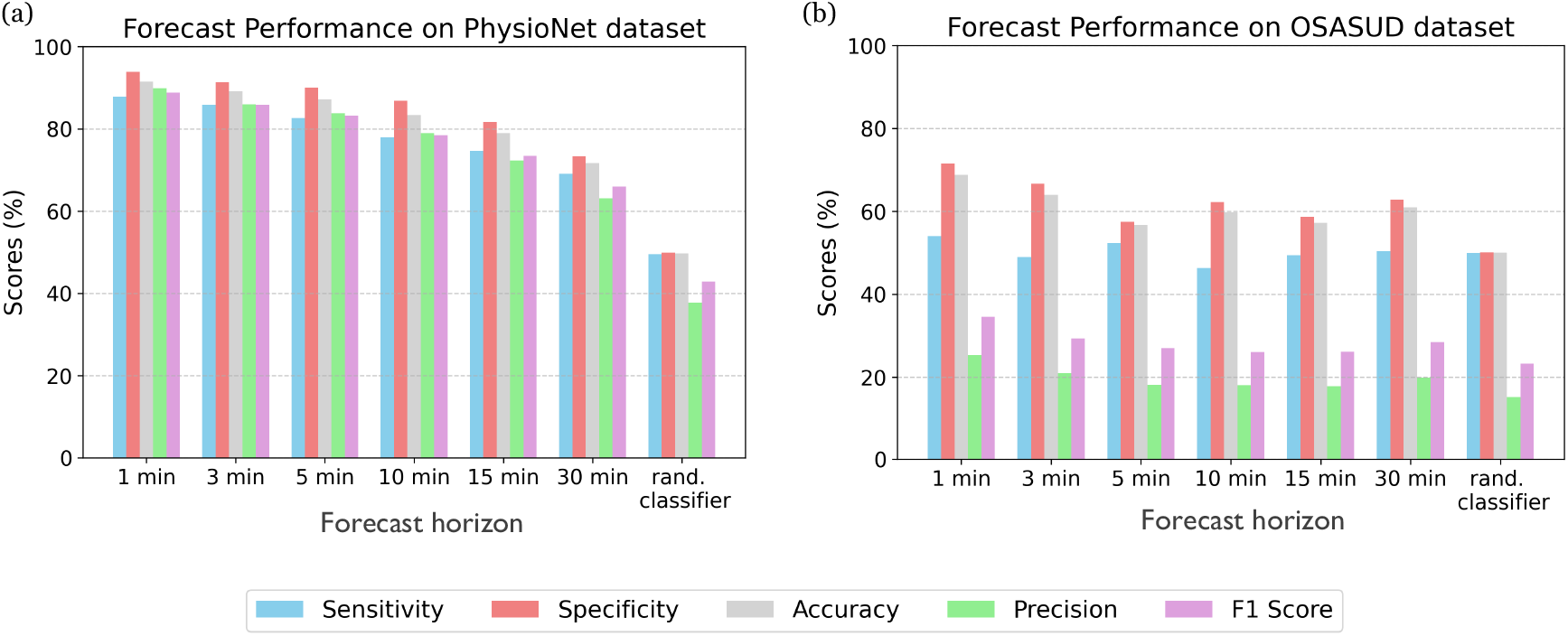
Forecast performance on the PhysioNet Apnea-ECG and OSASUD datasets. We evaluate the forecast performance of our transformer model on the (**a**) PhysioNet Apnea-ECG Database and the (**b**) OSASUD dataset for different forecast horizons. For both datasets, the performance degrades as the forecast horizon increases, indicating the challenge in making predictions further into the future.

##### OSA Severity Diagnosis

We evaluate the ApneaFormer’s ability to diagnose the severity of apnea on the PhysioNet Apnea-ECG dataset. For each test patient, the Apnea Hypopnea Index (AHI) value is computed based on the predicted per-minute apnea labels. Following the classification scheme from Hu et al. (2022)^29^, a patient is classified as normal if AHI *<* 5, mild if 5 ≤AHI *<* 15, moderate if 15 ≤AHI *<* 30, and severe if AHI ≥ 30. In Fig. 6 (a), we present the confusion matrix for the true and predicted apnea severity categories, illustrating the model’s performance across different severity levels. In Fig. 6 (b), we show the raw diagnostic accuracy for each severity category. The model demonstrates perfect diagnostic accuracy (100%) for the normal and moderate categories, while there are slight misclassifications in the mild and severe categories.

**Figure 6.**
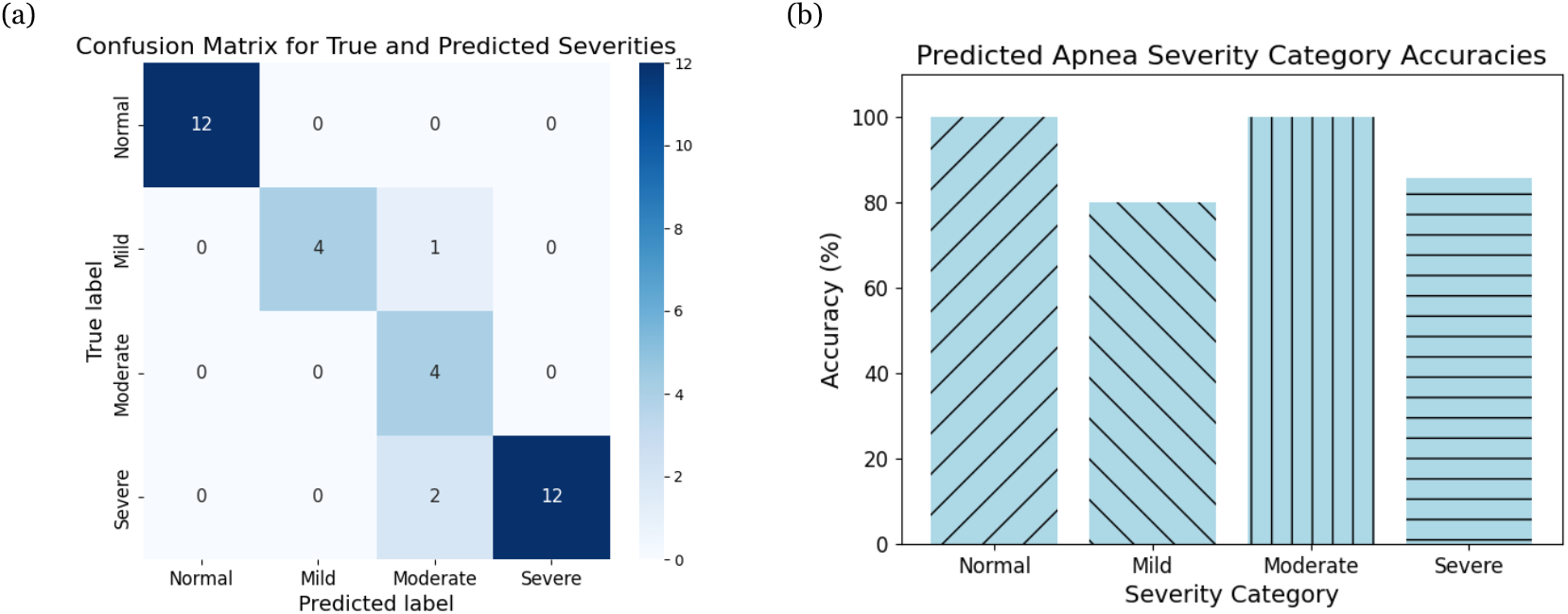
Diagnostic performance on PhysioNet Apnea-ECG dataset. We assess the diagnostic performance of our ApneaFormer model on the PhysioNet Apnea-ECG Database via (**a**) confusion matrix of true and predicted diagnostic categories based on the minutely apnea hypopnea index (AHI) and (**b**) the raw diagnostic accuracy by category.

##### Transfer Learning Experiments

Transfer learning allows a model trained on one dataset to be adapted to use with another, helping with generalization. In this study, we applied transfer learning in two ways. First, we trained a model on the PhysioNet Apnea-ECG dataset, which includes a large number of patients, and then finetuned the model for the smaller St. Vincent’s dataset. Second, we explored a patient-specific approach where a model was initially trained on a subset of patients from the same dataset (Vincent dataset) and then fine-tuned on individual patient during testing. This aligns with the principles of precision medicine, aiming to tailor the model to individual patient characteristics.

###### (a) Transfer Learning across datasets

The primary motivation for transfer learning in this context is to leverage a large source dataset to improve model performance on a smaller target dataset. In this study, we use the MIT PhysioNet Apnea-ECG dataset, which contains 70 patients, as the source dataset and the St. Vincent’s University Hospital dataset, comprising 25 patients, as the target dataset. Pretraining the model on PhysioNet Apnea-ECG allows it to learn from a larger patient cohort, enhancing its generalization capability. Fine-tuning on the training data from the St. Vincent’s dataset further adapts the model to region-specific patterns influenced by geographic, dietary and lifestyle factors unique to the patient population.

We experimented with training on varying numbers of patients from the target dataset to study the impact of dataset size on model performance. The F1 score as a function of training size is shown in Fig. 7 (a). Across all training set sizes, models utilizing transfer learning (TL) outperform those trained solely on the target dataset from random initialization, as indicated by the consistently higher F1 scores of TL-based model (red curve) compared to the baseline (blue curve). Additionally, we investigated the effect of fine-tuning different subsets of the model parameters. Specifically, we experimented with freezing varying numbers of convolutional (Conv) blocks during fine-tuning. The results indicate that fine-tuning all model parameters (red curve) yields the best overall performance, particularly for larger training sizes. However, for limited training data, fine-tuning fewer parameters leads to better F1 scores, suggesting that partial-fine tuning avoids the risk of overfitting while still enabling knowledge transfer. Notably, as the training size increases, all models—including those without transfer learning—exhibit an improvement in F1 score. However, the gap between the TL-based models and the baseline remains significant, showing the advantage of leveraging pretrained knowledge from a larger source dataset.

**Figure 7.**
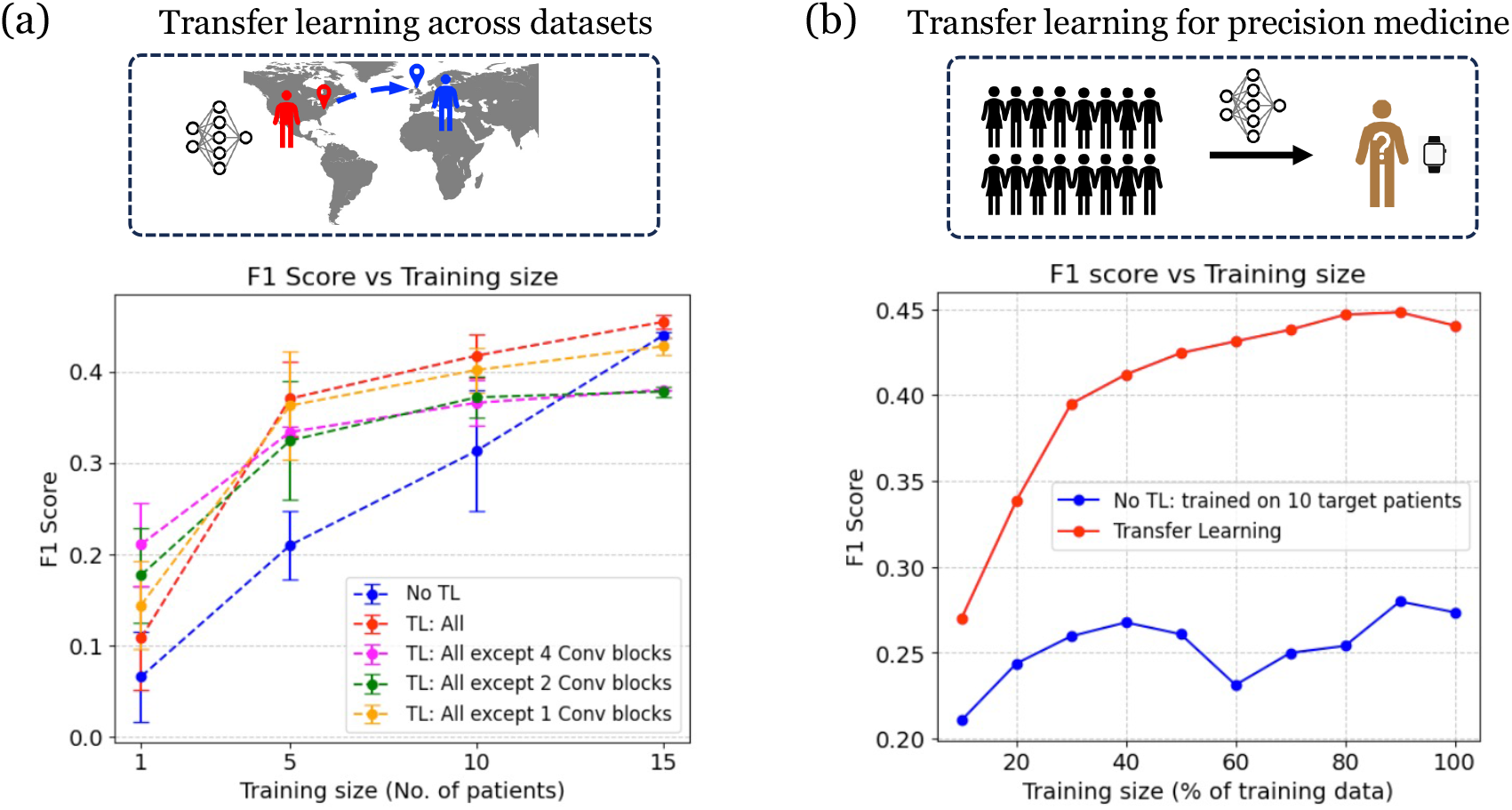
F1 score as a function of training data for transfer learning experiments. In (a) the PhysioNet Apnea-ECG dataset is used as the source dataset and the Vincent dataset is used as the target dataset. In (b), transfer learning is used to improve patient-wise personalized predictions. Here, the first 15 patients from the Vincent dataset are used as the source dataset, and partial data of each of the remaining 10 patients is used as the target dataset.

###### (b) Transfer learning for personalized predictions

The motivation for this experiment is to study how transfer learning can improve model predictions for individual patients by leveraging prior knowledge from a larger cohort. To achieve this, we first pretrain a source model using data from 15 patients in the St. Vincent dataset. For each of the remaining 10 patients, the last 20% of their data is reserved for testing, while varying fractions of the first 80% are used for training. For each patient, the source model is fine-tuned on the available training data and then evaluated on the corresponding test data. This process is repeated for all patients, and the micro-averaged evaluation metrics are computed. We conduct this experiment across different training sizes, and the results are shown in Fig. 7 (b). The red curve represents the performance of the model with transfer learning, while the blue curve corresponds to training from scratch without transfer learning (i.e., using only the target patient’s data). The results show that transfer learning consistently outperforms training from scratch across all training sizes. Notably, even with a small fraction of training data, the transfer learning model achieves significantly higher F1 scores. As the training size increases, the performance gap remains substantial, highlighting the benefits of transfer learning in improving personalized predictions.

## DISCUSSION

In this study, we propose and validate a holistic framework for obstructive sleep apnea (OSA) detection, severity classification, and forecasting using electrocardiogram (ECG) signals. By leveraging a combination of feature engineering rooted in dynamical systems theory, statistical signal processing, and modern deep learning architectures, our approach offers a scalable and interpretable solution to a complex clinical problem traditionally dependent on resource-intensive polysomnography.

Our work builds upon the physiological link between apnea events and alterations in cardiac activity, as reflected in ECG signals. Through feature engineering techniques inspired by dynamical systems — such as Lyapunov exponent and kurtosis — alongside classical statistical descriptors, we extract interpretable features that capture both nonlinear and linear dynamics of the cardiac signal during apneic events. Importantly, this allows our models to not only achieve strong predictive performance, but also provide physiologically meaningful insight into the underlying signal alterations associated with OSA.

We systematically evaluated both conventional machine learning models and modern deep learning architectures on multiple datasets of increasing complexity. While classical machine learning models achieved reasonable performance on clean datasets, their accuracy degraded significantly when applied to the more clinically complex OSASUD dataset. In contrast, the ApneaFormer, our deep learning model demonstrated robust performance across both datasets. A key innovation of ApneaFormer is its ability to process the full-resolution ECG signal without aggressive downsampling, thereby preserving fine-grained temporal features critical for detecting the subtle physiologic changes that accompany apnea.

Beyond static detection, we extended our framework to two clinically relevant tasks: forecasting and severity classification. The ability to forecast upcoming apnea events introduces potential applications in anticipatory interventions or real-time therapeutic adjustments. Similarly, by aggregating per-minute model predictions, we derived patient-level apnea-hypopnea indices (AHI), enabling severity classification that aligns with standard clinical diagnostic thresholds. This framework can also be adapted to predict hypoxia by incorporating real-time data from wearable biometric devices, which could enable early detection of hypoxic episodes and facilitate timely therapeutic interventions.

Transfer learning emerged as a particularly powerful component of our framework, addressing one of the key challenges in clinical machine learning: generalization across diverse patient populations. We demonstrated that models pretrained on large datasets can be effectively adapted to smaller datasets using cross-dataset transfer learning, improving performance in limited-data settings. Furthermore, we explored patient-specific fine-tuning, showing that personalized models further improve prediction accuracy, even when patient-level training data are sparse. These findings underscore the potential of transfer learning to facilitate precision medicine approaches in sleep apnea management.

Despite these advances, several limitations should be acknowledged. While our framework incorporates physiologically inspired features such as RR interval, Lyapunov exponent, and kurtosis, it remains primarily data-driven and does not explicitly model the causal physiological pathways underlying OSA. Future work could incorporate more formal mechanistic or causal modeling to better capture the complex cardiorespiratory interactions involved in apnea. In addition, the inclusion of complementary physiologic signals — such as oxygen saturation (SpO2), respiratory effort, or multimodal wearable sensor data — may further enhance model performance, particularly for long-range forecasting tasks where prediction accuracy declines as the forecast horizon increases.

In summary, our work represents a significant step forward in the application of artificial intelligence to sleep medicine. The ability to non-invasively detect, classify, and forecast OSA from ECG signals — a widely available biometric which may be obtained from wearables — offers a pathway to more continuous, scalable, and patient-centered diagnostic and monitoring solutions. As AI continues to evolve, such frameworks may play an increasingly central role in expanding access to diagnostic tools, enabling earlier interventions, and personalizing care for patients with OSA and related cardiorespiratory conditions.

## METHODS

### Dataset

Data was collected from three separate databases, the PhysioNet Apnea-ECG Database^11^, the Obstructive Sleep Apnea Stroke Unit Dataset (OSASUD)^8^, and the St. Vincent’s University Hospital / University College Dublin Sleep Apnea Database^34^. The PhysioNet Apnea-ECG Database is comprised of 70 nighttime ECG recordings. The ECG recordings have a sampling rate of 100 Hz and are single channel, which is consistent among all patients in the database. Each minute of the ECG is annotated as either apneic or non-apneic breathing, and this scoring is based on standard polysomnographic methods. The recordings range from 401 – 578 minutes in length. Data in the PhysioNet Apnea-ECG Database was collected from two studies: one in which the effects of obstructive sleep apnea on arterial blood pressure were assessed in patients with moderate to severe sleep apnea, and another in which data was collected from both healthy volunteers and volunteers with sleep apnea. Thirty-five patients are divided into a training subset, which contains a total of 17,125 labelled minutes, and the remaining patients are divided into a test subset, which contains a total of 17,303 labelled minutes^11^.

The OSASUD dataset is comprised of polysomnographic records from 30 patients who were admitted to the stroke unit of the Udine University Hospital in Italy. The ECG recordings have a sampling rate of 80 Hz, and data is collected for 12 leads. ECG data is publicly available for three leads, leads I, II, and III, and data processing is done for lead I. Each second of data is annotated as either apneic or non-apneic breathing; this assessment was made by a trained physician based on standard polysomnographic methods. The recordings range from 246 – 714 minutes in length. The patients in the OSASUD dataset are not divided into a training and test subset. Additionally, the authors indicate that this dataset is more representative of real-world patient data than competing datasets, which collect recordings in idealized circumstances and have strict exclusion criteria (Bernardini et al., 2022).

The St. Vincent’s University Hospital / University College Dublin Sleep Apnea Database is comprised of 25 overnight polysomnographic records from adult patients suspected to have disordered breathing during sleep. The ECG recordings are obtained from a simultaneous three-channel Holter ECG (V5, CC5, and V5R) at 128 Hz, and each minute of disordered breathing is labeled as either apneic or non-apneic breathing. The labeling was performed by a sleep technologist. The recordings range from 355 - 462 minutes in length, and the patients are not divided into a training or test subset^34^.

### Data Processing

#### Conventional ML Data Processing

For use in conventional machine learning models, ECG preprocessing was applied identically to the PhysioNet Apnea-ECG and OSASUD datasets to ensure consistency. ECG signals were denoised using NeuroKit2^47^ by applying a fifth-order high-pass Butterworth filter (0.5 Hz) to remove low-frequency noise and a powerline filter at 50 Hz to remove powerline interference.

Afterwards, OSASUD ECG signals, originally sampled at 80 Hz, were resampled to 100 Hz via linear interpolation to match the PhysioNet sampling rate. R peaks were detected using a gradient-based method, which first located QRS complexes using the steepness of the ECG signal’s absolute gradient and idenified R-peaks with the local minima of the gradient during the QRS complexes. The amplitude of the ECG at the R-peak locations was then stored as the R-amplitude. The interval between adjacent R-peaks (RR interval) was then calculated by subtracting the time between adjacent R-peaks, which was determined using the indexes of the R-peaks. Finally, the first-order difference of the RR intervals was determined by taking the difference between two adjacent RR intervals. Therefore, the four basic features include the denoised ECG, the R amplitude, the RR intervals, and the RR interval first-order difference, forming four core features following Hu et al.^29^. For the PhysioNet Apnea-ECG Database, minutely apnea labels were assigned to each ECG datapoint in its corresponding minute, and for the OSASUD dataset, secondly apnea labels were assigned to each ECG datapoint in its corresponding second.

Morphological features, including the mean, skewness, and kurtosis, were computed per second (100 data points), and 10-second rolling means of these features were calculated to capture local trends. To prevent data loss during the final ten seconds of a recording, rolling means were calculated with the maximum number of remaining data points.

Afterwards, NeuroKit2^47^ was used to automatically obtain heart rate variability (HRV) features. These features were calculated for 60-second windows of the denoised ECG and are derived from analyses in the time domain, frequency domain, and nonlinear domain. Time-domain-derived features, which are difference-based and deviation-based features calculated from the RR interval, provide basic information about a patient’s HRV. Frequency-domain-derived features analyze the distribution of absolute or relative power in different frequency bands (such as low frequency (LF), high frequency (HF), and the LF/HF ratio), providing valuable information about the relation between HRV and autonomic behavior. Finally, nonlinear-domain-derived features, including Poincaré plot geometries, entropy measurements, and fluctuation analyses, capture complex dynamics and patterns in heart activity that are not evident in linear analyses^48^. For several patients in both datasets, NeuroKit2 was unable to extract HRV features for specific segments of the ECG. In these cases, the portion of ECG that could not undergo feature extraction was removed.

To reduce the size of the data, the sampling rate was lowered from 100 Hz to 3 Hz by linear interpolation. Finally, static covariates were appended, including patient age, sex, height, and weight for the PhysioNet Apnea-ECG Database and patient age, sex, and body mass index (BMI) for the OSASUD dataset. While such features are not attainable through the ECG, they provide easily attainable and valuable biometric information about the patient. Data was stored on a patient-by-patient basis in Pandas DataFrames^49^.

Finally, thresholding of select features was performed. Specifically, ECG amplitudes and R amplitudes were clipped to a minimum and maximum of −5 and 5 mV, in accordance with the ordinary limits for ECG amplitudes^50^. For the OSASUD dataset, the ECG and R amplitudes were scaled by 0.1 to align with the PhysioNet Apnea-ECG Database. Then, the RR interval and RR interval first-order difference were clipped to a maximum of 2 seconds, which was selected based on a study determining the maximum RR interval to be approximately 2 seconds in a study collecting 24-hour ECG recordings of 1,528 patients^51^.

#### Deep Learning Data Processing

The ECG data in the PhysioNet Apnea-ECG dataset were acquired at a sampling frequency of 100 Hz. A second-order Butterworth bandpass filter was applied to the signal, with a low-pass cutoff of 5 Hz and a high-pass cutoff of 35 Hz. To address outliers, ECG values for each patient were clipped to a range defined by the mean ± 10 standard deviations. Subsequently, each patient’s ECG signal were normalized to the [0,1] range. The filtered signals were then segmented into overlapping 3-minute windows (18,000 data points), with the label for each segment indicating whether the middle minute of the window corresponded to an apnea event (1 data point). These 3-minute windows served as the input to the deep learning models, and the corresponding label represented the target output.

For the OSASUD dataset, we used the ECG signal from lead I, which was sampled at 80 Hz. Each patient’s ECG signal was clipped within the range of 100 to 200 units before normalizing to the [0, 1] range. Similar to the PhysioNet Apnea-ECG dataset, the filtered signals were segmented into 3-minute windows (14,400 data points). The apnea annotations in the OSASUD dataset were provided at a 1-second resolution, as opposed to the 1-minute resolution in the PhysioNet Apnea-ECG dataset. Therefore, each segment is associated with 60 labels, indicating whether each second in the middle minute corresponds to an apnea event (60 data points). Missing ECG data points were imputed with −1, and segments with more than 50 % missing values were excluded from the train-test data.

For the Vincent dataset, the ECG signal was acquired at 128 Hz frequency. The ECG signal was filtered using the *ecg_ clean* method of the NeuroKit2 Python library^47^. The filtered ECG signal was normalized to the [0,1] range and resampled to 100 Hz. Similar to the other datasets, the filtered signal was segmented into 3-minutes (18,000 data points). Each segment was associated with 60 labels indicating the annotation for each second of the middle minute.

In the case of PhysioNet Apnea-ECG and OSASUD datasets, we additionally experimented with three signals for the deep learning models: i) RR interval, ii) Lyapunov exponent, and iii) kurtosis, each computed at a frequency of 1 Hz (180 data points for each 3-minute window). The RR interval is computed after cleaning the normalized ECG signal using the *ecg_clean* method from the NeuroKit2 Python library. RR intervals were calculated as the distance between the R-peaks, identified using the *ecg_peaks* method of the NeuroKit2 library. The RR intervals were clipped at 2 s and then scaled by a factor of 0.5 to normalize the values to [0,1] range. The RR intervals were then resampled to a 1 Hz frequency, such that each 3-minute window corresponds to 180 data points of RR intervals. The Lyapunov exponent was calculated using Rosenstein’s algorithm (Rosenstein et al., 1993)^52^ for one-minute long sliding windows of the ECG signal for both the PhysioNet Apnea-ECG and the OSASUD datasets. The calculation was performed using the Nolds (NOnLinear measures for Dynamical Systems) Python package (Schölzel, 2019)^53^. For the PhysioNet Apnea-ECG dataset, we used an embedding embedding dimension of 10, a *τ* of 1/50, a minimum separation of 100 time steps, and a trajectory length of 100 time steps with a fit offset of 20 time steps were used to calculate the Lyapunov exponent. The ECG signal was downsampled from 100 Hz to 50 Hz prior to Lyapunov exponent calculation. For the OSASUD dataset, we used an embedding dimension of 15, *τ* of 1/20, a minimum seperation of 40 time steps, a trajectory length of 50 time steps and a fit offset of 20 time steps were used. The ECG signal was downsampled from 80 Hz to 20 Hz prior to the Lyapunov exponent calculation for the OSASUD dataset. Similar to the Lyapunov exponents, the kurtosis was calculated for one minute sliding window. The kurtosis was calculated using the *kurtosis* method in scipy.stats Python library.

### Correlational Studies

An exploratory data analysis was performed on both the PhysioNet Apnea-ECG Database and the OSASUD dataset. NeuroKit2-derived HRV features that had missing entries were dropped. Summary statistics were obtained for the entire dataset, as well as for subsets according to apnea severity and apnea diagnosis. Secondly, statistical analyzes were performed to identify significant differences between the severity groups of apnea and between healthy patients and patients diagnosed with apnea. Mann-Whitney U tests were used to assess differences in medians and distributions of features between patients diagnosed with apnea and patients not diagnosed with apnea. Assumptions of the Mann-Whitney U test were verified. Specifically, the single dependent variable of each test is continuous, the independent variable is categorical with two groups, the observations are independent, and the distributions for the groups of the independent variable are of similar shape. A level of significance of 0.05 was used.

### Explainable AI

The SHAP (SHapley Additive exPlanations) method is a technique proposed by Lundberg and Lee (2017)^33^to explain the decisions made by machine learning models. This method is based on game theory and calculates the marginal contribution of each feature to a model prediction using *Shapley* values. In other words, the difference between a model’s prediction for a specific instance and the average prediction across the entire dataset is distributed among the features according to the principle of additive contribution. The SHAP value calculated for each feature quantitatively indicates to what extent that feature influenced the prediction in a positive or negative direction. In this way, the outputs of complex models, often referred to as “black boxes,” become more interpretable. This is particularly valuable in the healthcare domain, where it helps reveal which indicators the model relied on to reach a decision, thus providing high interpretability.

The SHAP method uses the following formula to compute the contribution of each feature to the model prediction:

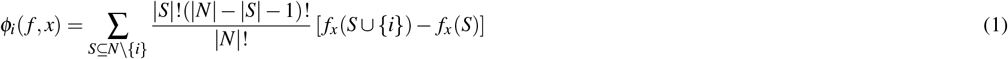

where *N* denotes the set of all features, and *f*_*x*_(*S*) represents the model output conditioned on subset *S*. SHAP adopts efficient approximation algorithms, such as TreeSHAP, to compute these values in practice. According to Lundberg and Lee^33^, SHAP provides desirable theoretical properties, including local accuracy and consistency.

### Model Architectures

#### Conventional ML Models

In this study, conventional machine learning models were employed to detect obstructive sleep apnea (OSA) using ECG-derived features from the PhysioNet Apnea-ECG and OSASUD datasets. The models evaluated include Logistic Regression (LR), Extreme Gradient Boosting (XGBoost), and Random Forest (RF). These models were selected due to their robustness, interpretability, and widespread use in classification tasks involving biomedical data.

**Logistic Regression (LR)** was utilized as a baseline model due to its simplicity and interpretability. It models the probability of an apnea event based on a linear combination of input features, making it suitable for assessing the statistical significance of ECG-derived features. LR was applied with L2 regularization to mitigate overfitting, and class weights were adjusted to address the imbalanced nature of the apnea and non-apnea labels.

**Extreme Gradient Boosting (XGBoost)**^54^, a tree-based ensemble method, was employed to capture complex, non-linear relationships within the ECG data. XGBoost leverages gradient boosting to iteratively improve model performance by minimizing a loss function. Hyperparameters, such as learning rate, maximum tree depth, and number of estimators, were tuned using grid search to optimize performance. The model’s ability to handle imbalanced datasets was enhanced through scale-positive-weight adjustments.

**Random Forest (RF)**^55^, another ensemble method, was used to provide robustness against noise and overfitting. By constructing multiple decision trees and aggregating their predictions, RF effectively captures feature interactions. The model was configured with a tuned number of trees and maximum depth, and feature importance was assessed to identify key ECG-derived predictors of apnea.

To ensure robust and reproducible model training, we performed systematic hyperparameter tuning for all conventional machine learning models evaluated in this study. For LR, we employed an elastic net penalty with a regularization strength (C) of 0.1, a solver set to ‘saga’, and an L1 ratio of 0, effectively emphasizing L2 regularization within the elastic net framework. For the XGBoost model, key parameters included a learning rate of 0.1, 1000 estimators, and a maximum tree depth of 5, enabling the model to capture complex patterns while maintaining generalizability. The RF classifier was configured with 1000 trees and no limit on tree depth (max_depth=None), allowing each tree to fully grow and capture hierarchical data relationships. These hyperparameter settings were determined via grid search and applied consistently throughout the experimental evaluations reported in Table 1.

The performance of these models was evaluated across four feature sets: (1) **Three Features** (R amplitude, RR interval, and first-order difference of RR interval), (2) **All Features** (including static covariates), (3) **Significant Features** (features showing statistical differences between apnea and non-apnea groups), and (4) **Significant ECG Features** (statistically significant features derived solely from ECG data). Performance metrics, including sensitivity, specificity, accuracy, precision, and F1 score, were computed to assess model efficacy (see Table 1).

Statistical significance of the conventional ML model features was determined using Mann-Whitney U tests with a level of significance of 0.05 to compare distributions between patients diagnosed with apnea and patients without apnea in the PhysioNet Apnea-ECG and the OSASUD datasets. Features found to be significant in the PhysioNet dataset were used to construct the **Significant Features** and **Significant ECG Features** sets, as this dataset offered a larger sample size and more reliable ECG signal quality compared to the OSASUD dataset, which exhibited limitations related to clinical variability, noise, and sample size.

#### Deep Learning Models

Two transformer models were presented in this work; the Hybrid Transformer model, and the ApneaFormer, shown in Fig. 8. The Hybrid Transformer model was designed to detect sleep apnea from ECG-derived time series features at a one-minute resolution. The developed architecture was inspired by the works of Hu et al.^29^ and Li et al. (2023), involving multiple representation pathways to capture temporal features. The model combines dilated, parallel, and depth-wise separable convolutions to extract multi-scale features from the input, the details of which are shown in Fig. 8 (a). These features are then concatenated and passed through a squeeze-and-excitation block^56^ to adaptively recalibrate the channel-wise feature sets. Following the squeeze-and-excitation block, the output is fed into a linear layer, and positional encoding^57^ is applied to retain sequence information. The encoded sequence is then passed through three transformer encoders, enabling the model to capture both local and long-range dependencies in the features. Lastly, the output is mapped to apnea or non-apnea logits via a linear projection.

**Figure 8.**
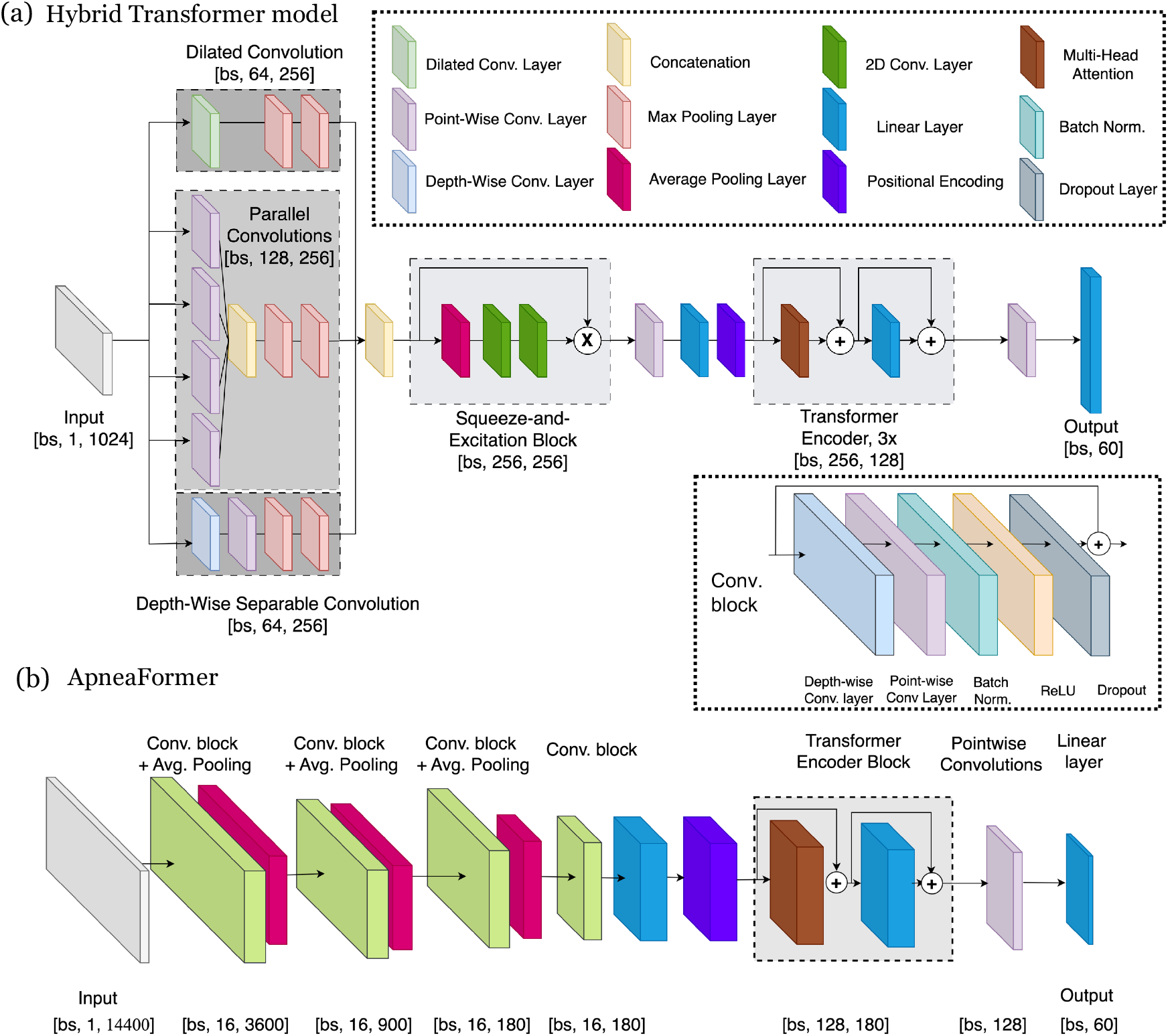
Architecture of the proposed deep learning models. Illustration of the model architecture for (a) Hybrid transformer model and (b) ApneaFormer model. Both the models consist of two key components: a convolutional stack and a transformer encoder block. The provided shapes of the output tensors of various layers are for the case of OSASUD dataset.

The ApneaFormer model is designed to process the ECG signal at its original resolution, allowing it to retain the full temporal information of the input data. Adapted from the AIOSA model proposed by Bernardini et al. (2021)^43^, the ApneaFormer consists of two primary components: a convolutional stack and a transformer encoder block, as shown in Fig. 8 (b). The input to the model is a 3-minute long ECG signal, represented by 14,400 time steps in the OSASUD dataset (sampled at 80 Hz). The sequence of convolutional blocks and average pooling helps to extract a learned representation of the original ECG signal with a lower temporal resolution while increasing the number of features/channels. Each convolutional block consists of depthwise separable convolutions with a filter size of 16 and kernel size of 3, followed by pointwise convolutions, batch normalization, and a ReLU activation function. Average pooling is applied at the end of first three convolutional blocks to reduce the temporal dimension, with kernel sizes of 4, 4, and 5 for OSASUD, and 4, 5, and 5 for PhysioNet. After processing through these blocks, the intermediate output is a tensor of shape [bs, 16, 180], denoting the learned compressed representation of the original 3-minute ECG signal. The output from the convolutional stack is processed using a transformer encoder block. The transformer encoder block is key to learning the temporal dependencies across the 180 time steps. We used the sinusoidal positional encoding prior to the encoder block. The output of the transformer encoder is a tensor of shape [bs, 128, 180] where 180 represents the number of times steps and 128 represents the number of features. The 180 feature vectors are combined into a single 128 dimensional feature vector using pointwise convolution, and subsequently passed through a linear layer to produced 60 output logits - one for each second in the middle minute of the input segment. The ApneaFormer model builds on the AIOSA model^43^ with important modifications for performance enhancement. Unlike AIOSA, which uses a bidirectional LSTM to process sequential data, ApneaFormer incorporates a transformer encoder block, enabling the model to learn long range temporal dependencies more effectively. In addition, ApneaFormer differs from AIOSA in how it handles the sequence of output vectors from biLSTM block. While AIOSA selects only the last output vector from the LSTM sequence, ApneaFormer takes into account all output vectors by performing a linear combination using a point-wise 1D convolution. This approach allows the model to capture patterns across the entire sequence rather than relying solely on the final representation.

In the PhysioNet Apnea-ECG dataset, the input has 18,000 time steps instead of the 14,400 time steps of the OSASUD dataset due to a higher sampling frequency of 100 Hz. Here, the output linear layer maps the features to a one-dimensional space since the apnea annotation are at a 1-minute resolution instead of a 1-second resolution. In the case of using additional features such as RR interval, Lyapunov exponent, or kurtosis are used, an additional transformer encoder block is used as described in the Supplementary Materials. The outputs of the ECG and auxiliary branches are then fused before classification. All models were trained using a weighted squared hinge loss to address class imbalance. For the OSASUD dataset, models were trained with a batch size of 256 for 200 epochs. For the PhysioNet dataset, we used a batch size of 512 and trained for 300 epochs. Optimization was performed using the Adam optimizer with a one-cycle learning rate scheduler. Final predictions were obtained by thresholding tte output logits at zero.

### Model Evaluations

The described models were evaluated for their performance on sleep apnea detection. The PhysioNet Apnea-ECG database and St. Vincent’s University Hospital / University College Dublin Sleep Apnea Database were evaluated using minutely classification of sleep apnea, and the OSASUD dataset was evaluated using secondly classification of sleep apnea. The evaluated performance metrics included sensitivity, specificity, accuracy, precision, and F1 score, where *TP* is the number of true positives, *FP* is the number of false positives, *TN* is the number of true negatives, and *FN* is the number of false negatives.

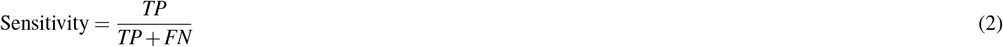

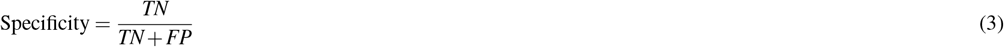

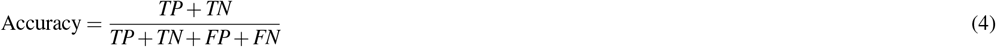

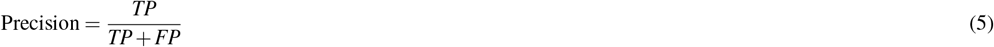

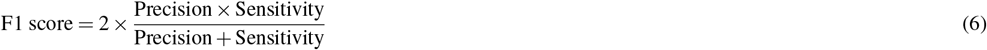

The transformer-based models were also evaluated using receiver operating characteristics (ROC), which plots the true positive rate (sensitivity) against the false positive rate (specificity). The models were compared to a random classifier, which has an area under the curve (AUC) of 0.5, and areas greater than 0.5 indicate better performance than would be attained by random classification.

Finally, transformer models were evaluated for their performance on sleep apnea diagnosis using a similar approach to Hu et al. (2022)^29^. The Apnea Hypopnea Index (AHI) for each patient in the test sets were calculated as follows:

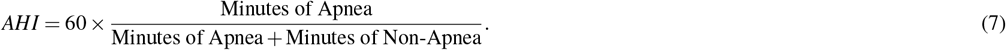

## DATA AVAILABILITY

We used publicly available datasets in this study. The PhysioNet Apnea-ECG dataset^11^ is publicly accessible and can be downloaded from https://physionet.org/content/apnea-ecg/1.0.0/. The OSASUD dataset^8^ can be downloaded from https://springernature.figshare.com/collections/A_dataset_of_stroke_unit_recordings_for_the_detection_of_Obstructive_Sleep_Apnea_Syndrome/5630890/1. The St. Vincent dataset can be dowloaded from https://physionet.org/content/ucddb/1.0.0/.

## CODE AVAILABILITY

The code for our work is available at the GitHub repository: https://github.com/alanjohnvarghese/ApneaFormer.

## ACKNOWLEDGEMENTS

The authors thank Andrei Petrus for his prior work on the detection of sleep apnea using machine learning algorithms in the Obstructive Sleep Apnea Stroke Unit Dataset. This work was partially supported by a grant from KardioStatus (Grant No: GR5253063).

## AUTHOR CONTRIBUTIONS

G.E.K. supervised the work. G.E.K., A.C. and A.P. formulated the problem. A.J.V., A.G. and M.A. developed the model. A.J.V., A.G. and M.A. implemented the computer code and performed computations. A.J.V., A.G., M.A. and V.O. analyzed data. A.J.V., A.G., M.A., V.O., A.P., A.C. and G.E.K. wrote the paper. A.J.V. and A.G. contributed equally to this work.

## References

1. Epstein, L. J. et al. Clinical guideline for the evaluation, management and long-term care of obstructive sleep apnea in adults. J. clinical sleep medicine 5, 263–276 (2009).

2. Lévy, P. et al. Obstructive sleep apnoea syndrome. Nat. Rev. Dis. Primers 1, 15015, DOI: 10.1038/nrdp.2015.15 (2015).

3. Motamedi, K. K., McClary, A. C. & Amedee, R. G. Obstructive sleep apnea: a growing problem. Ochsner J. 9, 149–153 (2009).

4. Zheng, N. S. et al. Sleep patterns and risk of chronic disease as measured by long-term monitoring with commercial wearable devices in the all of us research program. Nat. Medicine 30, 2648–2656, DOI: 10.1038/s41591-024-03155-8 (2024).

5. Mazzotti, D. R. & Manetta, M. Uncovering the role of sleep on human health. Nat. Medicine 31, 737–738, DOI: 10.1038/s41591-025-03529-6 (2025).

6. Jean-Louis, G., Zizi, F., Clark, L. T., Brown, C. D. & McFarlane, S. I. Obstructive sleep apnea and cardiovascular disease: role of the metabolic syndrome and its components. J. Clin. Sleep Medicine 4, 261–272 (2008).

7. Wang, Y. et al. Association between sleep duration and hypertension risk in patients with obstructive sleep apnea. npj Prim. Care Respir. Medicine 35, 26, DOI: 10.1038/s41533-025-00429-7 (2025).

8. Bernardini, A., Brunello, A., Gigli, G. L., Montanari, A. & Saccomanno, N. OSASUD: A dataset of stroke unit recordings for the detection of Obstructive Sleep Apnea Syndrome. Sci. Data 9, 177 (2022).

9. Retamales, G. et al. Towards automatic home-based sleep apnea estimation using deep learning. npj Digit. Medicine 7, 144, DOI: 10.1038/s41746-024-01139-z (2024).

10. Park, J. H., Wang, C. & Shin, H. FDA-cleared home sleep apnea testing devices. npj Digit. Medicine 7, 123, DOI: 10.1038/s41746-024-01112-w (2024).

11. Penzel, T., Moody, G. B., Mark, R. G., Goldberger, A. L. & Peter, J. H. The apnea-ECG database. In Computers in Cardiology 2000. Vol. 27 (Cat. 00CH37163), 255–258 (IEEE, 2000).

12. Gerstenslager, B. & Slowik, J. M. Sleep study. In StatPearls (StatPearls Publishing, Treasure Island (FL), 2023). Updated 2023 Aug 14.

13. Byun, J.-H., Kim, K. T., Moon, H.-j., Motamedi, G. K. & Cho, Y. W. The first night effect during polysomnography, and patients’ estimates of sleep quality. Psychiatry research 274, 27–29 (2019).

14. Tamaki, M., Nittono, H., Hayashi, M. & Hori, T. Examination of the first-night effect during the sleep-onset period. Sleep 28, 195–202 (2005).

15. Mincholé, A. & Rodriguez, B. Artificial intelligence for the electrocardiogram. Nat. Medicine 25, 22–23, DOI: 10.1038/s41591-018-0306-1 (2019).

16. Hannun, A. Y. et al. Cardiologist-level arrhythmia detection and classification in ambulatory electrocardiograms using a deep neural network. Nat. Medicine 25, 65–69, DOI: 10.1038/s41591-018-0268-3 (2019).

17. Thomas, R. J., Mietus, J. E.Peng, C.-K. & Goldberger, A. L. An electrocardiogram-based technique to assess cardiopulmonary coupling during sleep. Sleep 28, 1151–1161 (2005).

18. Bradicich, M. et al. Nocturnal heart rate variability in obstructive sleep apnoea: a cross-sectional analysis of the sleep heart health study. J. thoracic disease 12, S129 (2020).

19. Hassan, A. R. Computer-aided obstructive sleep apnea detection using normal inverse Gaussian parameters and adaptive boosting. Biomed. Signal Process. Control. 29, 22–30 (2016).

20. Hassan, A. R. & Haque, M. A. Computer-aided obstructive sleep apnea identification using statistical features in the EMD domain and extreme learning machine. Biomed. Phys. & Eng. Express 2, 035003 (2016).

21. Sharma, H. & Sharma, K. An algorithm for sleep apnea detection from single-lead ECG using Hermite basis functions. Comput. biology medicine 77, 116–124 (2016).

22. Song, C., Liu, K., Zhang, X., Chen, L. & Xian, X. An obstructive sleep apnea detection approach using a discriminative hidden Markov model from ECG signals. IEEE Trans. on Biomed. Eng. 63, 1532–1542 (2015).

23. Martin-Gonzalez, S. et al. Heart rate variability feature selection in the presence of sleep apnea: An expert system for the characterization and detection of the disorder. Comput. biology medicine 91, 47–58 (2017).

24. Sharma, H. & Sharma, K. Sleep apnea detection from ECG using variational mode decomposition. Biomed. Phys. & Eng. Express 6, 015026 (2020).

25. Bahrami, M. & Forouzanfar, M. Detection of sleep apnea from single-lead ECG: Comparison of deep learning algorithms. In 2021 IEEE International Symposium on Medical Measurements and Applications (MeMeA), 1–5 (IEEE, 2021).

26. Qin, H. & Liu, G. A dual-model deep learning method for sleep apnea detection based on representation learning and temporal dependence. Neurocomputing 473, 24–36 (2022).

27. Faust, O., Barika, R., Shenfield, A., Ciaccio, E. J. & Acharya, U. R. Accurate detection of sleep apnea with long short-term memory network based on RR interval signals. Knowledge-Based Syst. 212, 106591 (2021).

28. Shao, S. et al. Obstructive sleep apnea detection scheme based on manually generated features and parallel heterogeneous deep learning model under IoMT. IEEE J. Biomed. Heal. Informatics 26, 5841–5850 (2022).

29. Hu, S., Cai, W., Gao, T. & Wang, M. A hybrid transformer model for obstructive sleep apnea detection based on self-attention mechanism using single-lead ECG. IEEE Trans. on Instrum. Meas. 71, 1–11 (2022).

30. Li, C. et al. TFformer: A time frequency information fusion based CNN-transformer model for OSA detection with single-lead ECG. IEEE Trans. on Instrum. Meas. (2023).

31. Zhang, H., Cisse, M., Dauphin, Y. N. & Lopez-Paz, D. mixup: Beyond empirical risk minimization. arXiv preprint 1710.09412 (2017).

32. Chawla, N. V., Bowyer, K. W., Hall, L. O. & Kegelmeyer, W. P. SMOTE: Synthetic Minority Over-sampling Technique. J. artificial intelligence research 16, 321–357 (2002).

33. Lundberg, S. M. & Lee, S.-I. A unified approach to interpreting model predictions. In Guyon, I. et al. (eds.) Advances in Neural Information Processing Systems, vol. 30 (Curran Associates, Inc., 2017).

34. Goldberger, A. et al. PhysioBank, PhysioToolkit, and PhysioNet: Components of a new research resource for complex physiologic signals. Circ. [Online] 101, 215–220 (2000).

35. Chattopadhyay, S., Sarkar, G. & Das, A. Sleep apnea diagnosis by DWT-based kurtosis, radar and histogram analysis of electrocardiogram. IETE J. Res. 66, 518–526 (2020).

36. Smbatovna, S. K. & Alekseevna, M. L. Recognition of sleep apnea by EEG using nonlinear dynamics methods. In 2021 Ural Symposium on Biomedical Engineering, Radioelectronics and Information Technology (USBEREIT), 0009–0011 (IEEE, 2021).

37. Varon, C., Caicedo, A., Testelmans, D., Buyse, B. & Van Huffel, S. A novel algorithm for the automatic detection of sleep apnea from single-lead ECG. IEEE transactions on biomedical engineering 62, 2269–2278 (2015).

38. Zhang, L., Fu, M., Xu, F., Hou, F. & Ma, Y. Heart rate dynamics in patients with obstructive sleep apnea: heart rate variability and entropy. Entropy 21, 927 (2019).

39. Ravelo-García, A. G. et al. Application of the permutation entropy over the heart rate variability for the improvement of electrocardiogram-based sleep breathing pause detection. Entropy 17, 914–927 (2015).

40. Pan, W.-Y. et al. Multiscale entropic assessment of autonomic dysfunction in patients with obstructive sleep apnea and therapeutic impact of continuous positive airway pressure treatment. Sleep Medicine 20, 12–17 (2016).

41. Han, Q. & Wang, P. Estimation of the largest Lyapunov exponent of the HRV signals. J. Biomed. Eng. 24, 732–735 (2007).

42. Tayel, M. B. & AlSaba, E. I. Robust and sensitive method of lyapunov exponent for heart rate variability. arXiv preprint 1508.00996 (2015).

43. Bernardini, A., Brunello, A., Gigli, G. L., Montanari, A. & Saccomanno, N. AIOSA: An approach to the automatic identification of obstructive sleep apnea events based on deep learning. Artif. Intell. Medicine 118, 102133 (2021).

44. Chang, H.-Y., Yeh, C.-Y.Lee, C.-T. & Lin, C.-C. A sleep apnea detection system based on a one-dimensional deep convolution neural network model using single-lead electrocardiogram. Sensors 20, 4157 (2020).

45. Shen, Q., Qin, H., Wei, K. & Liu, G. Multiscale deep neural network for obstructive sleep apnea detection using RR interval from single-lead ECG signal. IEEE Trans. on Instrum. Meas. 70, 1–13, DOI: 10.1109/TIM.2021.3062414 (2021).

46. Wang, T., Lu, C., Shen, G. & Hong, F. Sleep apnea detection from a single-lead ECG signal with automatic feature-extraction through a modified LeNet-5 convolutional neural network. PeerJ 7, e7731 (2019).

47. Makowski, D. et al. NeuroKit2: A python toolbox for neurophysiological signal processing. Behav. Res. Methods 53, 1689–1696, DOI: 10.3758/s13428-020-01516-y (2021).

48. Pham, T., Lau, Z. J., Chen, S. H. A. & Makowski, D. Heart rate variability in psychology: A review of HRV indices and an analysis tutorial. Sensors 21, DOI: 10.3390/s21123998 (2021).

49. Wes McKinney. Data Structures for Statistical Computing in Python. In Stéfan van der Walt & Jarrod Millman (eds.) Proceedings of the 9th Python in Science Conference, 56 – 61, DOI: 10.25080/Majora-92bf1922-00a (2010).

50. Xie, L., Li, Z., Zhou, Y., He, Y. & Zhu, J. Computational diagnostic techniques for electrocardiogram signal analysis. Sensors 20, DOI: 10.3390/s20216318 (2020).

51. Yanaga, T. et al. Usefulness of 24-hour recordings of electrocardiogram for the diagnosis and treatment of arrhythmias with special reference to the determination of indication of artificial cardiac pacing : Present status and future of clinical electrocardiology. Jpn. Circ. J. 45, 366–375, DOI: 10.1253/jcj.45.366 (1981).

52. Rosenstein, M. T., Collins, J. J. & De Luca, C. J. A practical method for calculating largest lyapunov exponents from small data sets. Phys. D: Nonlinear Phenom. 65, 117–134 (1993).

53. Schölzel, C. Nonlinear measures for dynamical systems (version 0.5. 2). zenodo (2019).

54. Chen, T. & Guestrin, C. XGBoost: A scalable tree boosting system. In Proceedings of the 22nd ACM SIGKDD international conference on knowledge discovery and data mining, 785–794 (2016).

55. Breiman, L. Random forests. Mach. learning 45, 5–32 (2001).

56. Hu, J., Shen, L. & Sun, G. Squeeze-and-excitation networks. In 2018 IEEE/CVF Conference on Computer Vision and Pattern Recognition, 7132–7141, DOI: 10.1109/CVPR.2018.00745 (2018).

57. Vaswani, A. et al. Attention is all you need. CoRR abs/1706.03762 (2017). 1706.03762.

