## Supplemental Materials for "A multifaceted approach for obstructive sleep apnea classification from ECG signal using deep learning"

### SUPPLEMENTARY MATERIALS

#### Abbreviations

We provide a list of key abbreviations used in Table 1.

| Abbreviation | Full Form |
|---|---|
| OSA | Obstructive Sleep Apnea |
| OSASUD | Obstructive Sleep Apnea Stroke Unit Dataset |
| SHAP analysis | SHapley Additive exPlanations analysis |
| KNN | K-Nearest Neighbors |
| BMI | Body Mass Index |
| HRV | Heart Rate Variability |
| HFD | Higuchi Fractal Dimension |
| LR | Logistic Regression |
| XGBoost | Extreme Gradient Boosting |
| RF | Random Forest |
| ROC | Receiver Operating Characteristic |
| AUC | Area Under the Curve |
| AHI | Apnea-Hypopnea Index |
| TL | Transfer Learning |
| Conventional ML | Conventional Machine Learning |
| SVM | Support Vector Machine |

**Table 1.** Table of abbreviations used in this study.

#### Statistically significant features

For all patients in the PhysioNet Apnea-ECG Database and the OSASUD dataset, mean values were calculated for each electrocardiogram-derived feature. First, features that had missing row entries in either dataset were dropped from both datasets. This resulted in the removal of 26 features, leaving a total of 79 features for the PhysioNet Apnea-ECG Database and 78 features for the OSASUD dataset. The discrepancy in feature counts between the datasets is due to the PhysioNet Apnea-ECG Database providing patient heights and weights, whereas the OSASUD dataset utilizes patient BMI instead. The mean values were then aggregated according to apnea diagnoses, where the two groups are patients with apnea and patients without apnea.

Table 2 presents the statistically significantly different features for patients with and without apnea in the PhysioNet Apnea-ECG Database. Multiple features were found to have statistically significant differences in the distributions of patients with apnea and patients without apnea. The R amplitude and RR interval, kurtosis of the ECG, and rolling mean of the kurtosis of the ECG were the four directly calculated features from the ECG that had statistically significant differences between patients with apnea and patients without apnea. Patient age, sex, and weight also had significantly different distributions between patients with apnea and patients without apnea, which is in agreement with research on sleep apnea risk factors[1–3]. A total of 42 HRV-associated NeuroKit2 features had significantly different distributions between patients with apnea and patients without apnea, indicating that HRV analysis provides a wealth of information regarding the differentiation of apnea and non-apnea.

Regarding the NeuroKit2-derived features, there were 12 significantly different time domain features, 4 significantly different frequency domain features, and 26 significantly different nonlinear domain features. Therefore, features in the nonlinear domain comprise a significant source of difference between patients with and without sleep apnea.

The same analysis was performed on the OSASUD dataset. Unlike the PhysioNet Apnea-ECG Database, two features, including HRV_pNN50 and BMI, rather than 49 were found to have statistically significant differences in the distributions of patients with apnea and patients without apnea. The discrepancy between the PhysioNet Apnea-ECG Database and the OSASUD dataset may have several origins, including the lower number of patients in the OSASUD dataset, differences in ECG recording data collection due to different clinical environments, inability to filter out noise adequately in the OSASUD dataset, or perhaps physiological differences.

#### Violin plots of Lyapunov exponent and kurtosis

In Fig. 1 we show the violin plots of Lyapunov exponent and kurtosis for patients in both PhysioNet Apnea-ECG and OSASUD datasets. The patients have been sorted based on AHI value to highlight potential trends in the violin plot related to the severity of apnea. In the PhysioNet Apnea-ECG dataset, the violin plot of patients with $AHI = 0$ have been masked for visual and interpretative clarity, as the variation in the distribution for these patients might arise from other underlying health conditions

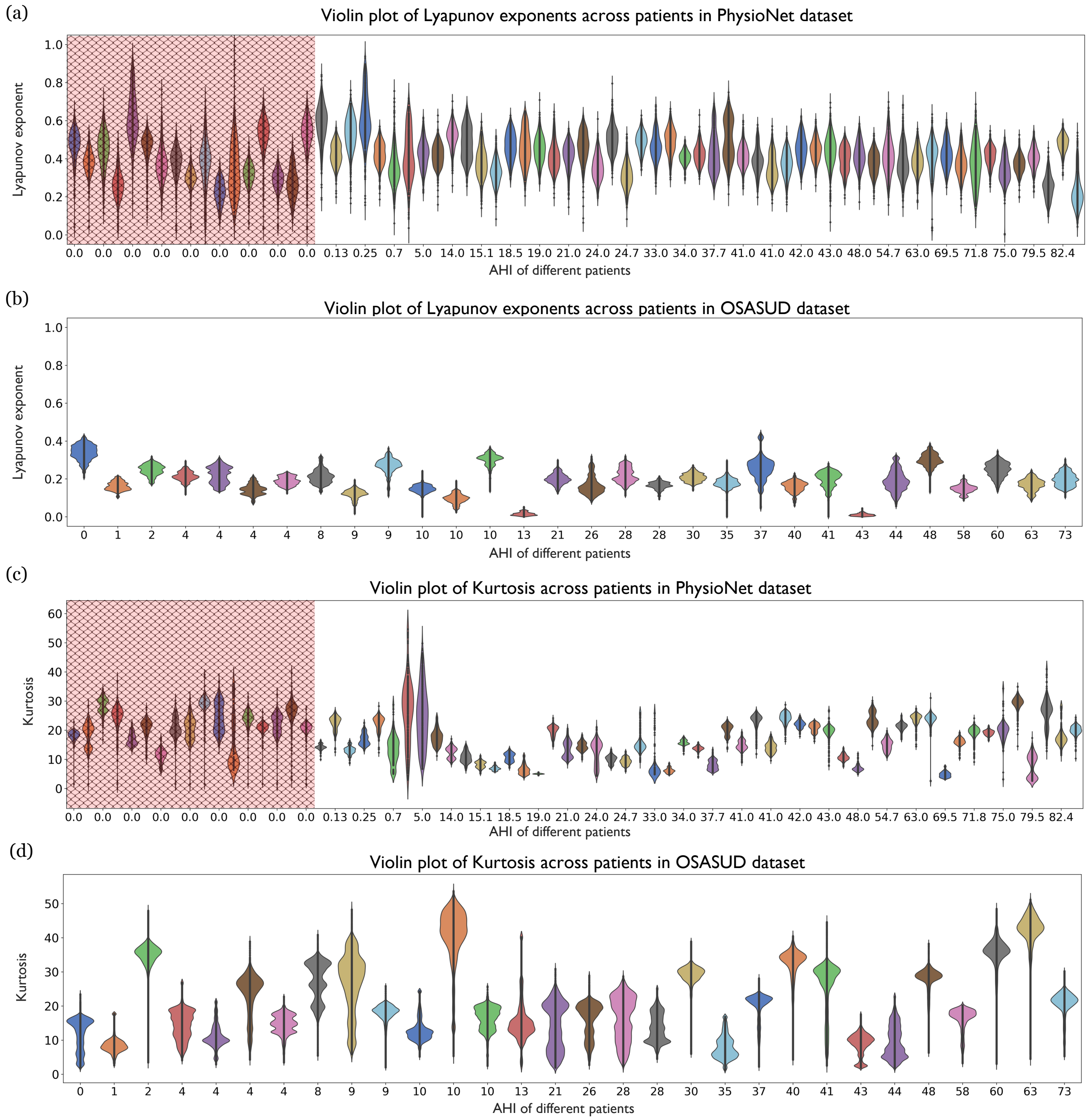


**Figure 1. Variation of Lyapunov exponent and kurtosis across patients in the PhysioNet Apnea-ECG and OSASUD datasets**. Violin plot of the Lyapunov exponent and kurtosis across patients in the (**a**) & (**c**) PhysioNet Apnea-ECG dataset and (**b**) & (**c**) OSASUD dataset. The patients have been sorted based on the AHI value, and is marked on the x-axis. We notice that as the severity of the apnea increases, the Lyapunov exponent has a decreasing trend across patients. The trends for the kurtosis is less clearer.

unrelated to apnea.. In the case of Lyapunov exponent, we see a slight downward shifting (decreasing) trend in the violin as AHI increases in both datasets. The trends are less pronounced in the OSASUD dataset potentially due to the presence of comorbidities. The violin plots for kurtosis does not reveal any clear trends in either dataset, suggesting that kurtosis may not be as sensitive to the severity of apnea.

#### Scalogram of ECG-derived features

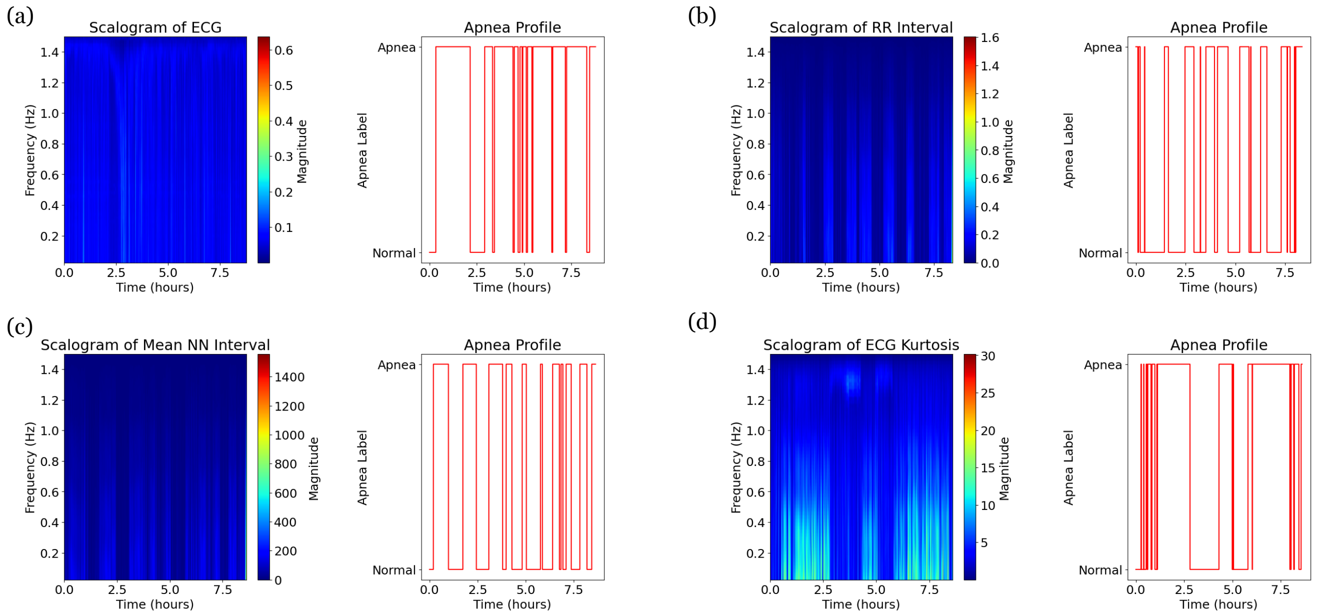


**Figure 2. Visualization of the scalogram of ECG and ECG-derived features.** We present representative scalograms coupled with patient apnea profiles for several ECG-derived features: (**a**) the denoised ECG signal, (**b**) the RR interval , (**c**) the mean NN interval, and (**d**) the kurtosis of the ECG.

In Fig. 2 we show the scalograms of ECG and ECG-derived features. We see the occurences of apnea correspond to regions with activity in the scalogram. Although not utilized in our work, scalograms can be used in computer vision based models to classify apnea along with the ECG signal in a multimodal framework.

#### Data Processing and Production of Electrocardiogram-Derived Scalograms

The same data preprocessing steps for conventional ML models were performed for the evaluation of apnea classification in the PhysioNet Apnea-ECG Database using all artificial intelligence algorithms. Patients were split into a training and a test set, as specified by the PhysioNet Apnea-ECG Database. Specifically, 35 patients are included in the training set, and 35 patients are included in the test set. Firstly, sex was coded as 0 for female patients and 1 for male patients. Thresholding of select features was performed as described in Chapter 2.2. Specifically, ECG and R amplitudes were clipped to a minimum and maximum of -5 and 5 mV[4], and the RR interval and RR interval first-order difference were clipped to a maximum of 2 seconds[5]. Then, NeuroKit2-derived HRV features that had missing entries were dropped.

Afterwards, time series data was normalized patient-by-patient using z-score normalization as shown in Equation 1:

$$x_{\text{norm}} = \frac{x - \mu}{\sigma} \tag{1}$$

where $x_{\text{norm}}$ is the normalized value of a feature at a point in time, $x$ is the value of a feature at the time point, $\mu$ is the mean value of the feature throughout the ECG recording, and $\sigma$ is the standard deviation of the feature throughout the ECG recording.

This normalization method scales the features such that they have a mean value of 0 and a standard deviation of 1. Static covariates, including patient age, height, and weight, were also normalized using z-score normalization. However, these features were normalized with respect to the means and standard deviations of these features for the entire training set, which was comprised of 35 patients as specified by the PhysioNet Apnea-ECG Database. With the goal of minutely apnea classification, 1 in 180 row values were selected for classification due to a sampling rate of 3 Hz. With this downscaled minutely sampling of the data, a total of 16,810 minutes is included in the training set, and a total of 17,143 minutes is included in the test set.

Scalograms were produced for several features derived from patient ECGs from the PhysioNet Apnea-ECG Database. ECG amplitudes, R amplitudes, RR intervals, and RR interval first-order differences were clipped according to the thresholds defined in Chapter 2.2. Then, PyWavelets (Lee et al., 2019) was used to produce scalograms of the denoised ECG, mean NN interval (average RR interval of detected R-peaks in a 60-second interval according to NeuroKit2) RR interval, and ECG kurtosis for each patient.

First, the continuous wavelet transform (CWT) was taken for the four aforementioned features to decompose them in the frequency and time domains. A complex Morlet wavelet with scales ranging from 1 to 63 was used. The equation for a complex Morlet wavelet is provided in Equation 2 (Lee et al., 2019):

$$\varphi(t) = \frac{1}{\sqrt{\pi B}} e^{-\frac{t^2}{2}} \cos(5t) e^{j2\pi Ct} \tag{2}$$

where $\varphi(t)$ is the complex Morlet wavelet as a function of time $t$, $B$ is the wavelet's bandwidth, and $C$ is the center frequency.

Then, for each of the four features, the Spearman correlation coefficient between the magnitude of the scalogram at specific scales and the occurrence of apnea was calculated. A total of six scales were sampled: the first scale (1.50 Hz), the second scale (0.75 Hz), the third scale (0.50 Hz), the fifth scale (0.30 Hz), the tenth scale (0.15 Hz), the fifteenth scale (0.10 Hz), and the thirtieth scale (0.05 Hz). The sampled scales and the corresponding frequencies were obatined using PyWavelets (Lee et al., 2019).

Average correlations were then calculated for patients in different apnea severity groups, as well as for healthy patients and patients diagnosed with apnea. Spearman correlation coefficients were also calculated between the raw features and the occurrence of apnea to serve as a comparison. Mann-Whitney U tests were used to assess differences in medians and distributions of the Spearman correlations between patients diagnosed with apnea and patients not diagnosed with apnea. Specifically, the single dependent variable of each test is continuous, the independent variable is categorical with two groups, the observations are independent, and the distributions for the groups of the independent variable are of similar shape. Assumptions were verified, and a level of significance of 0.05 was selected.

##### *Correlation between Scalograms and Apnea*

Scalograms for the denoised ECG, RR interval, mean NN interval, and ECG kurtosis were produced for the 70 recordings in the PhysioNet Apnea-ECG Database, and Spearman correlation coefficients were calculated between these raw features or specific frequency bands on their scalograms and the occurrence of apnea. Fig. 2 presents representative scalograms and apnea profiles for several patients for the denoised ECG, RR interval, mean NN interval, and kurtosis of the denoised ECG.

As seen in Fig. 2, time periods during which apnea occurs were reflected on the scalograms, particularly for the RR interval, mean NN interval, and ECG kurtosis. For the RR interval, mean NN interval, and ECG kurtosis, periods of apnea appeared to coincide with higher scalogram magnitudes. Therefore, scalogram-based analyses of an overnight ECG could provide a meaningful set of features for detection of sleep apnea. Whole scalograms could be utilized as image input or magnitudes could be extracted at specific scales, as done in this work.

Table 3 presents the mean Spearman correlation coefficient values for the four AHI severity groups, as well as for patients who are diagnosed with or without apnea. As the apnea severity increased, the correlation between the occurrence of apnea and the features increased, specifically for the RR interval, mean NN interval, and ECG kurtosis. For these features, it was also noted that the correlation is higher for patients diagnosed with apnea than for patients without apnea. The denoised ECG was not correlated with the occurrence of apnea when analyzed as a raw feature or when assessed for different scales of the scalogram; this finding agrees with the results of the Mann-Whitney U tests, which determined no significant difference between patients with apnea and patients without apnea for the denoised ECG correlations. Additionally, for the mean NN interval, the correlation increased as the scale increased, which corresponds to a decrease in frequency. A similar trend is also seen for the kurtosis of the ECG; the correlation became evident only at the tenth scale. Overall, the correlation between the occurrence of apnea and the selected features was highest for the mean NN interval for the thirtieth scale.

#### Transfer learning

##### *Transfer learning across datasets*

In this subsection, we provide a detailed description of our experimental setup for transfer learning across datasets. Transfer learning helps us to improve model performance on datasets with limited training data, as is the case with the St. Vincent dataset, which contains only 25 patients. By leveraging the larger PhysioNet Apnea-ECG dataset, which has more patients, we aim to improve the performance on St. Vincent dataset. We begin by training our model on the source dataset, PhysioNet Apnea-ECG, and then fine-tune it using the training patients (Patients 1-15) from the St. Vincent dataset. The fine-tuned model is then evaluated on the test patients (Patients 16-25) of the St. Vincent dataset. Micro-averaged performance metrics, including sensitivity, specificity, accuracy, precision and f1 score, are computed on the test dataset.

To explore the effect of varying training size (i.e., the number of patients used for finetuning), we randomly sample a fixed number of patients from the training pool to fine-tune the model. The fine-tuned model is subsequently evaluated on the test data. Given that the model can be sensitive to the specific patients sampled for finetuning, particularly when using few patients, we mitigate this sensitivity by repeating the sampling and finetuning process five times for each training size, and report the mean and standard deviation of the performance metrics.

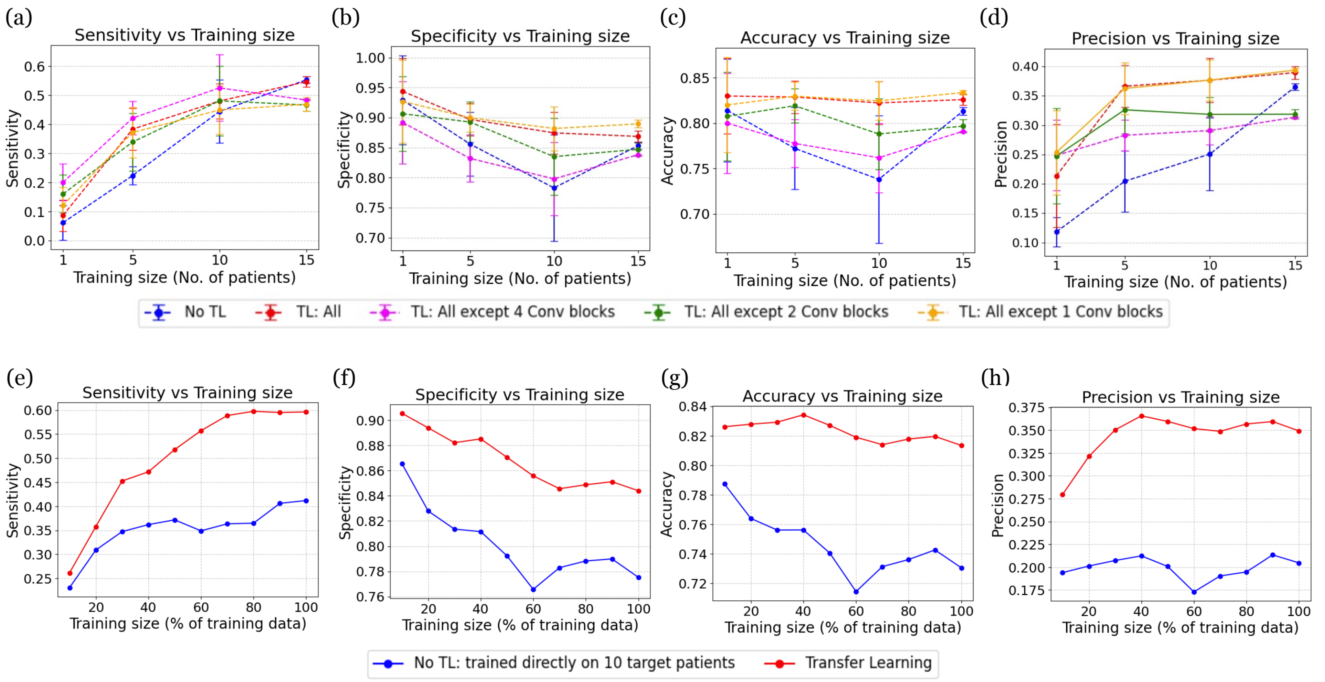


**Figure 3. Performance metrics for transfer learning experiments**. The top row (**a-d**) shows the sensitivity, specificity, accuracy and precision as a function of training size (number of patients) for transfer learning across datasets. The bottom row (**e-h**) shows the sensitivity, specificity, accuracy and precision as a function of training size (percentage of training data) for transfer learning to improve personalized predictions.

The setup described in the earlier paragraph is repeated for finetuning different parameter groups. The different curves in Fig. 3 correspond to tuning different parameters. The red curve corresponds to finetuning all the parameters. The yellow curve corresponds to finetuning all except the first convolutional block. The green curve corresponds to finetuning all except the first two convolutional blocks. The purple curve corresponds to finetuning all except 4 convolutional blocks. In essence, we progressively experiment with finetuning fewer and fewer parameters. We also compare the transfer learning experiments with baseline blue curve which corresponds to no transfer learning, that is, the model is trained from random initialization using the training data.

##### *Transfer learning for personalized predictions*

We utilized transfer learning to improve personalized predictions for the St. Vincent dataset. In this setup, Patients 1-15 served as the source dataset, while Patients 16-25 were used as the target dataset. Initially, the model was pretrained on the source dataset. For each patient in the target pool, the first 80% of the time series data was used to create 3-minute training windows, with the remaining 20% of the time-series reserved for testing. Micro-averaged evaluation metrics, including sensitivity, specificity, accuracy, precision and F1 score, were computed on the test data of the 10 target patients.

To analyse the effect of varying training sizes, we randomly sampled a fixed percentage of 3-minute windows for each patient in the target pool. The pre-trained model (trained on Patients 1-16) was finetuned using these sampled training data. The fine-tuned model was then evaluated on the corresponding patients test data. The micro-averaged evaluation metrics were reported across the 10 patients. This process was repeated for sampling percentages ranging from 10% to 100% in increments of 10%. The red curve in the second row of Fig. 3 shows the variation in evaluation metrics when transfer learning is applied. The blue curve represents the performance of the model without any transfer learning, where no source dataset is used for pretraining.

#### Model architecture when additional signals are used

In Fig. 6, we present the modified architecture of the ApneaFormer model when additional signals such as the RR interval, Lyapunov exponent, and kurtosis are included. These additional signals, typically available at a 1 Hz resolution, are represented as a sequence of length 180 for each 3-minute window. The additional signal is processed independently through a dedicated transformer encoder block, which first extracts temporal dependencies from the signal. The output of this encoder block is reduced to a single vector using pointwise convolution and then concatenated with the output vector from the block

**Algorithm 1** Transfer Learning across datasets. PhysioNet → Vincent Dataset

**Input:** Source dataset (PhysioNet) $\mathscr{D}_{\text{source}}$
**Input:** Target dataset (Vincent) with training pool $\mathscr{D}_{\text{train}} = \{P_1, P_2, \ldots, P_{15}\}$ and test data $\mathscr{D}_{\text{test}} = \{P_{16}, \ldots, P_{25}\}$, where each $P_i$ contains features $\mathbf{X}_i$ and labels $\mathbf{y}_i$
**Input:** Set of training sizes $\mathscr{K} = \{1, 5, 10, 15\}$ patients
**Input:** Learning algorithm $\mathscr{A}$ with hyperparameters $\theta$
**Input:** Number of repetitions $R = 5$
**Result:** Mean and std. of performance metrics for each training size

**Step 1:** Train base model $M_{\text{base}}$ on PhysioNet Apnea-ECG source dataset
$M_{\text{base}} \leftarrow \mathscr{A}(\mathscr{D}_{\text{source}}, \theta)$
**for** each $k \in \mathscr{K}$ **do** ▷ For each training size (1, 5, 10, or 15 patients)
  Initialize metrics storage across repetitions: $\mathscr{M}_k \leftarrow \emptyset$
  **for** $r = 1$ to $R$ **do** ▷ Step 5: Repeat 5 times
    **Step 2:** Randomly sample $k$ patients from $\mathscr{D}_{\text{train}}$: $\mathscr{S}_k \leftarrow \text{RandomSample}(\mathscr{D}_{\text{train}}, k)$
    **Step 3:** Fine-tune base model on sampled patients
    $M_{k,r} \leftarrow \text{Finetune}(M_{\text{base}}, \mathscr{S}_k, \theta)$
    **Step 4:** Evaluate fine-tuned model on test data and compute metrics
    Initialize: $TP \leftarrow 0, FP \leftarrow 0, TN \leftarrow 0, FN \leftarrow 0$
    **for** each $P_j \in \mathscr{D}_{\text{test}}$ **do**
      $\hat{\mathbf{y}}_j \leftarrow M_{k,r}(P_j)$ ▷ Generate predictions
      Calculate patient-level metrics $TP_j, FP_j, TN_j, FN_j$ using $\hat{\mathbf{y}}_j$ and $\mathbf{y}_j$
      $TP \leftarrow TP + TP_j$, $FP \leftarrow FP + FP_j$, $TN \leftarrow TN + TN_j$, $FN \leftarrow FN + FN_j$
    **end for**
    Compute micro-averaged performance metrics for this repetition:
    $\text{Sensitivity}_{k,r} \leftarrow \frac{TP}{TP+FN}$, $\text{Specificity}_{k,r} \leftarrow \frac{TN}{TN+FP}$, $\text{Accuracy}_{k,r} \leftarrow \frac{TP+TN}{TP+TN+FP+FN}$,
    $\text{Precision}_{k,r} \leftarrow \frac{TP}{TP+FP}$, $\text{F1}_{k,r} \leftarrow \frac{2\times TP}{2\times TP+FP+FN}$
    $\mathscr{M}_k \leftarrow \mathscr{M}_k \cup \{(\text{Sensitivity}_{k,r}, \text{Specificity}_{k,r}, \text{Accuracy}_{k,r}, \text{Precision}_{k,r}, \text{F1}_{k,r})\}$
  **end for**
  **Step 5:** Compute final statistics for training size $k$
    Calculate mean and standard deviation of each metric in $\mathscr{M}_k$ across the $R$ repetitions
**end for**
**Step 6:** Repeat entire process for all sample sizes in $\mathscr{K}$
**return** Mean and standard deviation of each metric for all training sizes

that processed the ECG signal. The concatenated features are subsequently passed through a series of linear layers, batch normalization, and dropout layers for final processing. This combined representation is used to make the final prediction, allowing the model to utilize both the ECG signal and the additional feature to classify apnea events.

When more than two additional features (consider $n$ features) are used at the same time, they are concatenated along the channel dimension to form a tensor of shape $[bs, n, 180]$. This tensor is first mapped to a tensor of shape $[bs, 128, 180]$ via a linear projection layer and then processed through a transformer encoder block as described above. This modified architecture jointly learns interactions between auxiliary signals and incorporate them alongside the ECG signal for improved apnea classification.

**Table 5.** Complete feature dictionary used in PhysioNet and OSASUD datasets. Feature definitions reproduced from NeuroKit2[6] under the MIT License.

| Feature | Datasets | Description |
|---|---|---|
| ECG | PhysioNet, OSASUD | Normalized ECG signal. |
| R_Amplitude | PhysioNet, OSASUD | Amplitude of R peak. |
| RR_Interval | PhysioNet, OSASUD | Time interval between R peaks. |
| RR_Interval_1D | PhysioNet, OSASUD | First-order difference of RR interval. |
| ECG_Mean | PhysioNet, OSASUD | Mean ECG signal (1 second). |
| ECG_Skew | PhysioNet, OSASUD | ECG skew (1 second). |
| ECG_Kurtosis | PhysioNet, OSASUD | ECG kurtosis (1 second). |

Continued on next page

| Feature | Datasets | Description |
|---|---|---|
| ECG_Mean_Rolling | PhysioNet, OSASUD | Rolling mean ECG signal (10 seconds). |
| ECG_Skew_Rolling | PhysioNet, OSASUD | Rolling ECG skew (10 seconds). |
| ECG_Kurtosis_Rolling | PhysioNet, OSASUD | Rolling ECG kurtosis (10 seconds). |
| Lyapunov_Exponent | PhysioNet, OSASUD | Calculated Lyapunov exponent (1 minute). |
| Lyapunov_Exponent_1D | PhysioNet, OSASUD | First-order difference of Lyapunov exponent. |
| 10-Minute_Kurtosis_1D | PhysioNet, OSASUD | First-order difference of 10-minute kurtosis. |
| HRV_MeanNN | PhysioNet, OSASUD | Mean of RR intervals (1 minute). |
| HRV_SDNN | PhysioNet, OSASUD | Standard deviation of RR intervals (1 minute). |
| HRV_RMSSD | PhysioNet, OSASUD | Square root of the mean of the squared differences of consecutive RR intervals (1 minute). |
| HRV_SDSD | PhysioNet, OSASUD | Standard deviation of the differences between RR intervals. |
| HRV_CVNN | PhysioNet, OSASUD | Standard deviation of SDNN divided by MeanNN (1 minute). |
| HRV_CVSD | PhysioNet, OSASUD | Root mean square of RMSSD divided by MeanNN (1 minute). |
| HRV_MedianNN | PhysioNet, OSASUD | Median of RR intervals (1 minute). |
| HRV_MadNN | PhysioNet, OSASUD | Median absolute deviation of RR intervals (1 minute). |
| HRV_MCVNN | PhysioNet, OSASUD | MadNN divided by MedianNN (1 minute). |
| HRV_IQRNN | PhysioNet, OSASUD | Intequartile range of RR intervals (1 minute). |
| HRV_SDRMSSD | PhysioNet, OSASUD | SDNN divided by RMSSD (1 minute). |
| HRV_Prc20NN | PhysioNet, OSASUD | 20th percentile of RR intervals (1 minute). |
| HRV_Prc80NN | PhysioNet, OSASUD | 80th percentile of RR intervals (1 minute). |
| HRV_pNN50 | PhysioNet, OSASUD | Percentage of absolute differences in consecutive RR intervals longer than 50 ms (1 minute). |
| HRV_pNN20 | PhysioNet, OSASUD | Percentage of absolute differences in consecutive RR intervals longer than 20 ms (1 minute). |
| HRV_MinNN | PhysioNet, OSASUD | Minimum RR interval (1 minute). |
| HRV_MaxNN | PhysioNet, OSASUD | Maximum RR interval (1 minute). |
| HRV_HTI | PhysioNet, OSASUD | HRV triangular index; total number of RR intervals divided by the height of the RR interval histogram (1 minute). |
| HRV_TINN | PhysioNet, OSASUD | Baseline width of RR intervals distribution obtained via triangular interpolation, where least squares error determines the triangle (1 minute). |
| HRV_HF | PhysioNet, OSASUD | Spectral power of high frequencies, 0.15 to 0.40 Hz (1 minute). |
| HRV_VHF | PhysioNet, OSASUD | Spectral power of very high frequencies, 0.40 to 0.50 Hz (1 minute). |
| HRV_TP | PhysioNet, OSASUD | Total spectral power (1 minute). |
| HRV_HFn | PhysioNet, OSASUD | Normalized high frequency; low frequency divided by total power (1 minute). |
| HRV_LnHF | PhysioNet, OSASUD | Log transformed HF (1 minute). |
| HRV_SD1 | PhysioNet, OSASUD | Standard deviation perpendicular to the Poincaré plot line of identity (1 minute). |
| HRV_SD2 | PhysioNet, OSASUD | Standard deviation along the Poincaré plot line of identity (1 minute). |
| HRV_SD1SD2 | PhysioNet, OSASUD | Ratio of SD1 t SD2 (1 minute). |
| HRV_S | PhysioNet, OSASUD | Area of ellipse described by SD1 and SD2 (1 minute). |
| HRV_CSI | PhysioNet, OSASUD | Cardiac symapthetic index; longitudinal variability of Poincaré plot divided by transverse variability (1 minute). |
| HRV_CVI | PhysioNet, OSASUD | Cardiac vagal index; logarithm of the product of longitudinal and transverse variability of the Poincaré plot (1 minute). |
| HRV_CSI_Modified | PhysioNet, OSASUD | Modified CSI; square of longitudinal variability divided by the transverse variability of the Poincaré plot (1 minute). |
| HRV_PIP | PhysioNet, OSASUD | Percentage of inflection points of the RR intervals (1 minute). |
| HRV_IALS | PhysioNet, OSASUD | Inverse of the average length of acceleration and deceleration segments (1 minute). |
| HRV_PSS | PhysioNet, OSASUD | Percentage of short segments (1 minute). |

Continued on next page

| Feature | Datasets | Description |
|---|---|---|
| HRV_PAS | PhysioNet, OSASUD | Percentage of NN intervals in alternation segments. |
| HRV_GI | PhysioNet, OSASUD | Guzik's index; in the Poincaré plot, distance of points above the line of identity to the line of identity divided by the distance of all points not on the line of identity to the line of identity (1 minute). |
| HRV_SI | PhysioNet, OSASUD | Slope index; in the Poincaré plot, phase angle of points above the line of identity divided by the phase angle of all points not on the line of identity (1 minute). |
| HRV_AI | PhysioNet, OSASUD | Area index; in the Poincaré plot, cumulative area of sectors corresponding to points above the line of identity divided by the cumulative area of sectors corresponding to all points not on the line of identity (1 minute). |
| HRV_PI | PhysioNet, OSASUD | Porta's index; in the Poincaré plot, number of points below the line of identity divided by the total number of points not on the line of identity (1 minute). |
| HRV_C1d | PhysioNet, OSASUD | Contributions of heart rate decelerations to the short-term HRV (1 minute). |
| HRV_C1a | PhysioNet, OSASUD | Contributions of heart rate accelerations to the short-term HRV (1 minute). |
| HRV_SD1d | PhysioNet, OSASUD | Short-term variance of the contributions of decelerations (1 minute). |
| HRV_SD1a | PhysioNet, OSASUD | Short-term variance of the contributions of accelerations (1 minute). |
| HRV_C2d | PhysioNet, OSASUD | Contributions of heart rate decelerations to the long-term HRV (1 minute). |
| HRV_C2a | PhysioNet, OSASUD | Contributions of heart rate accelerations to the long-term HRV (1 minute). |
| HRV_SD2d | PhysioNet, OSASUD | Long-term variance of the contributions of decelerations (1 minute). |
| HRV_SD2a | PhysioNet, OSASUD | Long-term variance of the contributions of accelerations (1 minute). |
| HRV_Cd | PhysioNet, OSASUD | Total contributions of heart rate decelerations to HRV (1 minute). |
| HRV_Ca | PhysioNet, OSASUD | Total contributions of heart rate accelerations to HRV (1 minute). |
| HRV_SDNNd | PhysioNet, OSASUD | Total variance of contributions of decelerations (1 minute). |
| HRV_SDNNa | PhysioNet, OSASUD | Total variance of contributions of accelerations (1 minute). |
| HRV_DFA_alpha1 | PhysioNet, OSASUD | Monofractal detrended fluctuation analysis of the heart rate signal for short-term correlations (1 minute). |
| HRV_MFDFA_alpha1_Width | PhysioNet, OSASUD | Multifractal detrended fluctuation analysis width of the singularity spectrum (1 minute). |
| HRV_MFDFA_alpha1_Peak | PhysioNet, OSASUD | Multifractal detrended fluctuation analysis value of singularity exponent H corresponding to the peak of singularity dimension D (1 minute). |
| HRV_MFDFA_alpha1_Mean | PhysioNet, OSASUD | Multifractal detrended fluctuation analysis mean of maximum and minimum values of singulariy exponent H (1 minute). |
| HRV_MFDFA_alpha1_Max | PhysioNet, OSASUD | Multifractal detrended fluctuation analysis value of singularity spectrum D corresponding to the maximum value of singularity exponent H (1 minute). |
| HRV_MFDFA_alpha1_Delta | PhysioNet, OSASUD | Multifractal detrended fluctuation analysis vertical distance between singularity spectrum D where the singularity exponents are at their minimum and maximum (1 minute). |
| HRV_MFDFA_alpha1_Asymmetry | PhysioNet, OSASUD | Multifractal detrended fluctuation analysis asymmetric ratio; centrality of the peak of the spectrum (1 minute). |
| HRV_MFDFA_alpha1_Fluctuation | PhysioNet, OSASUD | Multifractal detrended fluctuation analysis $h$-fluctuation index; power of the second derivative of $h$(q) (1 minute). |

Continued on next page

| Feature | Datasets | Description |
|---|---|---|
| HRV_MFDFA_alpha1_Increment | PhysioNet, OSASUD | Multifractal detrended fluctuation analysis cumulative function of the squared increments of the generalized Hurst's exponents between consecutive moment orders (1 minute). |
| HRV_ApEn | PhysioNet, OSASUD | Approximate entropy (1 minute). |
| HRV_ShanEn | PhysioNet, OSASUD | Shannon entropy (1 minute). |
| HRV_FuzzyEn | PhysioNet, OSASUD | Fuzzy entropy (1 minute). |
| HRV_MSEn | OSASUD | Multiscale entropy (1 minute). |
| HRV_CD | PhysioNet, OSASUD | Fractal correlation (1 minute). |
| HRV_HFD | PhysioNet, OSASUD | Higuchi fractal dimension (1 minute). |
| HRV_KFD | OSASUD | Katz's fractal dimension (1 minute). |
| HRV_LZC | PhysioNet, OSASUD | Lempel-Ziv complexity (1 minute). |
| Age | PhysioNet, OSASUD | Patient age. |
| Sex | PhysioNet, OSASUD | Patient sex. |
| Height | PhysioNet | Patient height. |
| Weight | PhysioNet | Patient weight. |
| BMI | OSASUD | Patient body mass index. |

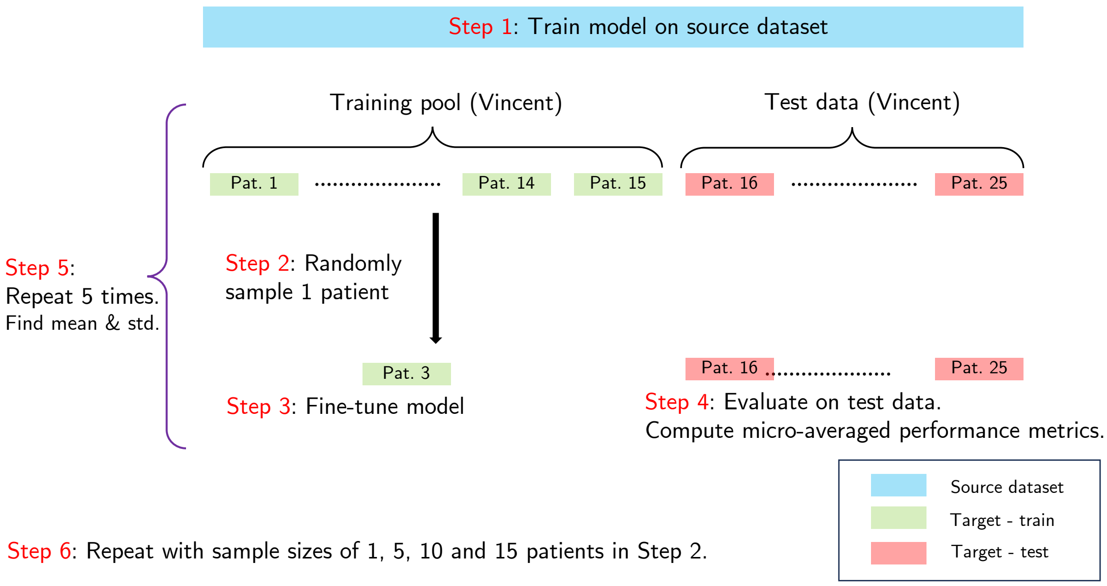


**Figure 4. Outline of the experimental setup for analyzing the effect of training size during transfer learning across datasets**. The model is first trained on PhysioNet Apnea-ECG dataset. The trained model is finetuned on varying numbers of randomly selected patients, and evaluated on a fixed pool of test patients.

---

**Algorithm 2** Transfer Learning for Personalized Predictions (Vincent dataset)

---

**Input:** Source dataset $\mathscr{D}_{\text{source}} = \{P_1, P_2, \ldots, P_{15}\}$ where each $P_i$ contains features $\mathbf{X}_i$ and labels $\mathbf{y}_i$
**Input:** Target pool $\mathscr{D}_{\text{target}} = \{P_{16}, \ldots, P_{25}\}$ where each $P_j$ contains features $\mathbf{X}_j$ and labels $\mathbf{y}_j$
**Input:** Set of fine-tuning fractions $\mathscr{F} = \{0.1, 0.2, \ldots, 1.0\}$ ▷ From 10% to 100%
**Input:** Learning algorithm $\mathscr{A}$ with hyperparameters $\theta$
**Result:** Performance metrics for each fine-tuning fraction

**Step 1:** Train base model $M_{\text{base}}$ on source dataset $\mathscr{D}_{\text{source}}$
$M_{\text{base}} \leftarrow \mathscr{A}(\mathscr{D}_{\text{source}}, \theta)$
**for each** $f \in \mathscr{F}$ **do** ▷ Step 6: Repeat for fraction varying from 10% to 100%
  Initialize: $TP \leftarrow 0$, $FP \leftarrow 0$, $TN \leftarrow 0$, $FN \leftarrow 0$
  **for each** $P_j \in \mathscr{D}_T$ **do** ▷ Step 4: Repeat for all patients in target pool
    **Step 2:** Split $P_j$ into fine-tuning set $P_j^{\text{train}}$ and test set $P_j^{\text{test}}$
      Sample a fraction $f$ from $P_j^{\text{train}}$ to get $P_{j,f}^{\text{train}}$.
    $M_j \leftarrow \text{Finetune}(M_{\text{base}}, P_{j,f}^{\text{train}}, \theta)$ ▷ Fine-tune model using fraction $f$ of patient data
    **Step 3:** Evaluate fine-tuned model on patient's test data
    $\hat{\mathbf{y}}_j \leftarrow M_j(P_j^{\text{test}})$ ▷ Generate predictions
    Calculate patient-level metrics $TP_j, FP_j, TN_j, FN_j$ using $\hat{\mathbf{y}}_j$ and $\mathbf{y}_j$
    $TP \leftarrow TP + TP_j$, $FP \leftarrow FP + FP_j$, $TN \leftarrow TN + TN_j$, $FN \leftarrow FN + FN_j$
  **end for**
  **Step 5:** Compute micro-averaged performance metrics for fraction $f$
  $\text{Sensitivity}_f \leftarrow \frac{TP}{TP+FN}$, $\text{Specificity}_f \leftarrow \frac{TN}{TN+FP}$, $\text{Accuracy}_f \leftarrow \frac{TP+TN}{TP+TN+FP+FN}$,
  $\text{Precision}_f \leftarrow \frac{TP}{TP+FP}$, $\text{F1}_f \leftarrow \frac{2\times TP}{2\times TP+FP+FN}$
**end for**
**return** Performance metrics for each fine-tuning fraction $f \in \mathscr{F}$

---

#### References


**1.** Franklin, K. A. & Lindberg, E. Obstructive sleep apnea is a common disorder in the population—a review on the epidemiology of sleep apnea. *J. Thorac. Dis.* **7** (2015).

**2.** Young, T., Evans, L., Finn, L. & Palta, M. Estimation of the clinically diagnosed proportion of sleep apnea syndrome in middle-aged men and women. *Sleep* **20**, 705–706, DOI: 10.1093/sleep/20.9.705 (1997). https://academic.oup.com/sleep/

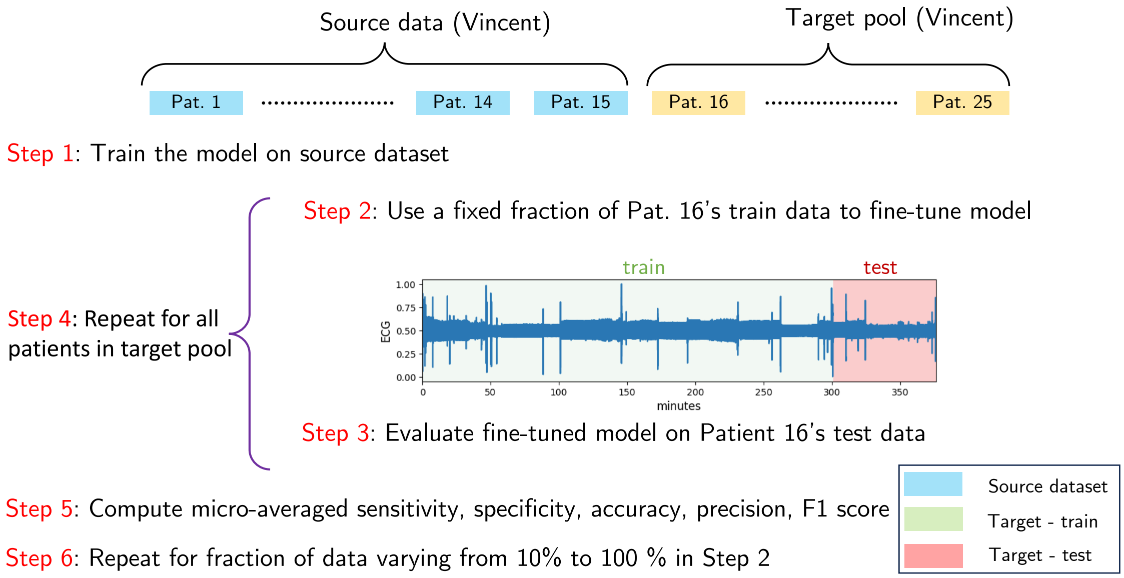


**Figure 5. Outline of experimental setup for analyzing the effect of training size for transfer learning to enhance personalized predictions.** The model is first trained on a fixed number of patients of St. Vincent dataset. The pretrained model is finetuned on a randomly sampled fraction of each test patient's data and evaluated on each patient's test data.

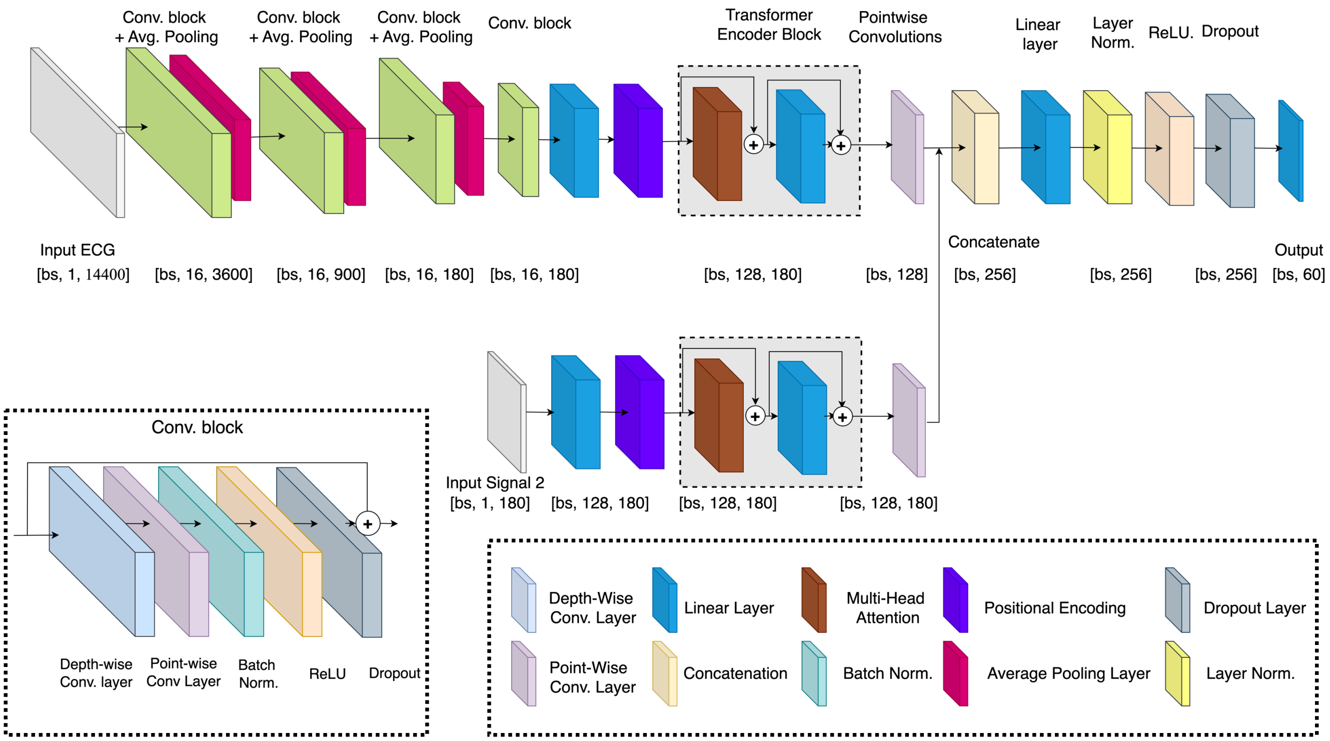


**Figure 6. ApneaFormer architecture when additional signals are used.** When input signals in addition to the ECG signal is used, they are fed in through a dedicated encoder block. The outputs from the two transformer blocks are fused together and processed further before obtaining the output logits.

article-pdf/20/9/705/25722419/sleep-20-9-705.pdf.

3. Randerath, W. *et al.* Challenges and perspectives in obstructive sleep apnoea. *Eur. Respir. J.* **52**, DOI: 10.1183/13993003.02616-2017 (2018).

4. Xie, L., Li, Z., Zhou, Y., He, Y. & Zhu, J. Computational diagnostic techniques for electrocardiogram signal analysis. *Sensors* **20**, DOI: 10.3390/s20216318 (2020).

5. Yanaga, T. *et al.* Usefulness of 24-hour recordings of electrocardiogram for the diagnosis and treatment of arrhythmias with special reference to the determination of indication of artificial cardiac pacing : Present status and future of clinical electrocardiology. *Jpn. Circ. J.* **45**, 366–375, DOI: 10.1253/jcj.45.366 (1981).

6. Makowski, D. *et al.* NeuroKit2: A python toolbox for neurophysiological signal processing. *Behav. Res. Methods* **53**, 1689–1696, DOI: 10.3758/s13428-020-01516-y (2021).

**Table 2. Statistically Significant Features.** Statistically significant features for the PhysioNet Apnea-ECG Database.

| Feature | Feature Subset |
|---|---|
| *Static Features* | |
| Age | — |
| Sex | — |
| Weight | — |
| *Non-NeuroKit2 Features* | |
| RR_Interval | — |
| R_Amplitude | — |
| ECG_Kurtosis | — |
| ECG_Kurtosis_Rolling | — |
| *Frequency Domain Features* | |
| HRV_HF | — |
| HRV_HFn | — |
| HRV_LnHF | — |
| HRV_TP | — |
| *Time Domain Features* | |
| HRV_pNN20 | — |
| HRV_SDRMSSD | — |
| HRV_pNN50 | — |
| HRV_MedianNN | — |
| HRV_Prc80NN | — |
| HRV_MeanNN | — |
| HRV_MaxNN | — |
| HRV_RMSSD | — |
| HRV_SDSD | — |
| HRV_CVSD | — |
| HRV_MinNN | — |
| HRV_Prc20NN | — |
| *Nonlinear Features* | |
| HRV_ApEn | Complexity/Fractal Physiology |
| HRV_CSI | Poincaré |
| HRV_DFA_alpha1 | Complexity/Fractal Physiology |
| HRV_FuzzyEn | Complexity/Fractal Physiology |
| HRV_LZC | Complexity/Fractal Physiology |
| HRV_MFDFA_alpha1_Max | Complexity/Fractal Physiology |
| HRV_PAS | Heart Rate Fragmentation |
| HRV_SD1SD2 | Poincaré |
| HRV_MFDFA_alpha1_Delta | Complexity/Fractal Physiology |
| HRV_CSI_Modified | Poincaré |
| HRV_Ca | Heart Rate Asymmetry |
| HRV_Cd | Heart Rate Asymmetry |
| HRV_S | Poincaré |
| HRV_SD1d | Heart Rate Asymmetry |
| HRV_CD | Complexity/Fractal Physiology |
| HRV_SD1 | Poincaré |
| HRV_SD1a | Heart Rate Asymmetry |
| HRV_C2a | Heart Rate Asymmetry |
| HRV_C2d | Heart Rate Asymmetry |
| HRV_C1a | Heart Rate Asymmetry |
| HRV_C1d | Heart Rate Asymmetry |
| HRV_CVI | Poincaré |
| HRV_MFDFA_alpha1_Width | Complexity/Fractal Physiology |
| HRV_PIP | Heart Rate Fragmentation |
| HRV_MFDFA_alpha1_Peak | Complexity/Fractal Physiology |
| HRV_PI | Heart Rate Asymmetry |

**Table 3. Spearman Correlation Summary.** Mean Spearman correlation coefficients between the occurrence of apnea and the denoised ECG, RR interval, mean NN interval, and kurtosis of the ECG for the 70 PhysioNet Apnea-ECG Database recordings.

| | Mean Values | | | | | | | |
|---|---|---|---|---|---|---|---|---|
| **Apnea Severity** | Correlation with ECG | Correlation @ Scale 1 | Correlation @ Scale 2 | Correlation @ Scale 3 | Correlation @ Scale 5 | Correlation @ Scale 10 | Correlation @ Scale 15 | Correlation @ Scale 30 |
| Normal (n = 23) | 0.002 | -0.003 | 0.000 | 0.002 | 0.003 | -0.001 | -0.002 | 0.004 |
| Mild (n = 5) | 0.009 | -0.049 | -0.072 | -0.066 | -0.041 | -0.024 | -0.023 | -0.025 |
| Moderate (n = 11) | -0.005 | 0.022 | 0.044 | 0.043 | 0.023 | 0.007 | 0.005 | 0.008 |
| Severe (n = 31) | 0.012 | -0.001 | -0.023 | -0.023 | -0.019 | -0.006 | 0.003 | 0.013 |
| **Apnea Diagnosis** | Correlation with ECG | Correlation @ Scale 1 | Correlation @ Scale 2 | Correlation @ Scale 3 | Correlation @ Scale 5 | Correlation @ Scale 10 | Correlation @ Scale 15 | Correlation @ Scale 30 |
| No Apnea (n = 23) | 0.002 | -0.003 | 0.000 | 0.002 | 0.003 | -0.001 | -0.002 | 0.004 |
| Has Apnea (n = 47) | 0.008 | -0.001 | -0.012 | -0.012 | -0.011 | -0.005 | 0.001 | 0.008 |
| p-value ($p < 0.05$) | - | - | - | - | - | - | - | - |
| **Apnea Severity** | Correlation with RR Interval | Correlation @ Scale 1 | Correlation @ Scale 2 | Correlation @ Scale 3 | Correlation @ Scale 5 | Correlation @ Scale 10 | Correlation @ Scale 15 | Correlation @ Scale 30 |
| Normal (n = 23) | -0.004 | 0.031 | 0.033 | 0.035 | 0.038 | 0.043 | 0.043 | 0.044 |
| Mild (n = 5) | 0.158 | 0.169 | 0.186 | 0.192 | 0.201 | 0.202 | 0.203 | 0.239 |
| Moderate (n = 11) | -0.047 | 0.231 | 0.249 | 0.259 | 0.270 | 0.245 | 0.246 | 0.286 |
| Severe (n = 31) | 0.231 | 0.230 | 0.244 | 0.251 | 0.252 | 0.204 | 0.164 | 0.224 |
| **Apnea Diagnosis** | Correlation with RR Interval | Correlation @ Scale 1 | Correlation @ Scale 2 | Correlation @ Scale 3 | Correlation @ Scale 5 | Correlation @ Scale 10 | Correlation @ Scale 15 | Correlation @ Scale 30 |
| No Apnea (n = 23) | -0.004 | 0.031 | 0.033 | 0.035 | 0.038 | 0.043 | 0.043 | 0.044 |
| Has Apnea (n = 47) | 0.158 | 0.224 | 0.239 | 0.247 | 0.251 | 0.213 | 0.188 | 0.240 |
| p-value ($p < 0.05$) | $p < 0.001$ | $p < 0.0001$ | $p < 0.0001$ | $p < 0.0001$ | $p < 0.0001$ | $p < 0.0001$ | $p < 0.0001$ | $p < 0.0001$ |
| **Apnea Severity** | Correlation with Mean NN Interval | Correlation @ Scale 1 | Correlation @ Scale 2 | Correlation @ Scale 3 | Correlation @ Scale 5 | Correlation @ Scale 10 | Correlation @ Scale 15 | Correlation @ Scale 30 |
| Normal (n = 23) | -0.019 | 0.003 | 0.006 | 0.004 | 0.002 | 0.017 | 0.023 | 0.032 |
| Mild (n = 5) | 0.172 | 0.116 | 0.128 | 0.132 | 0.130 | 0.180 | 0.193 | 0.267 |
| Moderate (n = 11) | -0.116 | 0.151 | 0.182 | 0.196 | 0.199 | 0.206 | 0.235 | 0.336 |
| Severe (n = 31) | 0.324 | 0.207 | 0.243 | 0.267 | 0.280 | 0.296 | 0.287 | 0.381 |
| **Apnea Diagnosis** | Correlation with Mean NN Interval | Correlation @ Scale 1 | Correlation @ Scale 2 | Correlation @ Scale 3 | Correlation @ Scale 5 | Correlation @ Scale 10 | Correlation @ Scale 15 | Correlation @ Scale 30 |
| No Apnea (n = 23) | -0.019 | 0.003 | 0.006 | 0.004 | 0.002 | 0.017 | 0.023 | 0.032 |
| Has Apnea (n = 47) | 0.205 | 0.184 | 0.217 | 0.236 | 0.245 | 0.262 | 0.265 | 0.359 |
| p-value ($p < 0.05$) | $p < 0.01$ | $p < 0.0001$ | $p < 0.0001$ | $p < 0.0001$ | $p < 0.0001$ | $p < 0.0001$ | $p < 0.0001$ | $p < 0.0001$ |
| **Apnea Severity** | Correlation with ECG Kurtosis | Correlation @ Scale 1 | Correlation @ Scale 2 | Correlation @ Scale 3 | Correlation @ Scale 5 | Correlation @ Scale 10 | Correlation @ Scale 15 | Correlation @ Scale 30 |
| Normal (n = 23) | 0.001 | 0.007 | 0.007 | 0.006 | 0.005 | 0.010 | 0.013 | 0.023 |
| Mild (n = 5) | 0.092 | -0.040 | -0.044 | -0.042 | -0.001 | 0.045 | 0.093 | 0.145 |
| Moderate (n = 11) | 0.043 | 0.053 | 0.062 | 0.060 | 0.048 | 0.056 | 0.081 | 0.182 |
| Severe (n = 31) | 0.106 | -0.039 | -0.045 | -0.042 | 0.008 | 0.122 | 0.135 | 0.214 |
| **Apnea Diagnosis** | Correlation with ECG Kurtosis | Correlation @ Scale 1 | Correlation @ Scale 2 | Correlation @ Scale 3 | Correlation @ Scale 5 | Correlation @ Scale 10 | Correlation @ Scale 15 | Correlation @ Scale 30 |
| No Apnea (n = 23) | 0.001 | 0.007 | 0.007 | 0.006 | 0.005 | 0.010 | 0.013 | 0.023 |
| Has Apnea (n = 47) | 0.090 | -0.017 | -0.020 | -0.018 | 0.016 | 0.098 | 0.118 | 0.199 |
| p-value ($p < 0.05$) | $p < 0.01$ | - | - | - | - | $p < 0.01$ | $p < 0.0001$ | $p < 0.0001$ |

**Table 4. Complete feature set.** List of all the features used in conventional ML models for PhysioNet Apnea-ECG dataset and OSASUD dataset.

| PhysioNet | OSASUD |
|---|---|
| ECG, R_Amplitude, RR_Interval, RR_Interval_1D, ECG_Mean, ECG_Skew, ECG_Kurtosis, ECG_Mean_Rolling, ECG_Skew_Rolling, ECG_Kurtosis_Rolling, Apnea, Lyapunov_Exponent, Lyapunov_Exponent_1D, 10-Minute_Kurtosis_1D, HRV_MeanNN, HRV_SDNN, HRV_RMSSD, HRV_SDSD, HRV_CVNN, HRV_CVSD, HRV_MedianNN, HRV_MadNN, HRV_MCVNN, HRV_IQRNN, HRV_SDRMSSD, HRV_Prc20NN, HRV_Prc80NN, HRV_pNN50, HRV_pNN20, HRV_MinNN, HRV_MaxNN, HRV_HTI, HRV_TINN, HRV_HF, HRV_VHF, HRV_TP, HRV_HFn, HRV_LnHF, HRV_SD1, HRV_SD2, HRV_SD1SD2, HRV_S, HRV_CSI, HRV_CVI, HRV_CSI_Modified, HRV_PIP, HRV_IALS, HRV_PSS, HRV_PAS, HRV_GI, HRV_SI, HRV_AI, HRV_PI, HRV_C1d, HRV_C1a, HRV_SD1d, HRV_SD1a, HRV_C2d, HRV_C2a, HRV_SD2d, HRV_SD2a, HRV_Cd, HRV_Ca, HRV_SDNNd, HRV_SDNNa, HRV_DFA_alpha1, HRV_MFDFA_alpha1_Width, HRV_MFDFA_alpha1_Peak, HRV_MFDFA_alpha1_Mean, HRV_MFDFA_alpha1_Max, HRV_MFDFA_alpha1_Delta, HRV_MFDFA_alpha1_Asymmetry, HRV_MFDFA_alpha1_Fluctuation, HRV_MFDFA_alpha1_Increment, HRV_ApEn, HRV_ShanEn, HRV_FuzzyEn, HRV_CD, HRV_HFD, HRV_LZC, Age, Sex, Height, Weight | ECG, R_Amplitude, RR_Interval, RR_Interval_1D, ECG_Mean, ECG_Skew, ECG_Kurtosis, ECG_Mean_Rolling, ECG_Skew_Rolling, ECG_Kurtosis_Rolling, Apnea, Lyapunov_Exponent, Lyapunov_Exponent_1D, 10-Minute_Kurtosis_1D, HRV_MeanNN, HRV_SDNN, HRV_RMSSD, HRV_SDSD, HRV_CVNN, HRV_CVSD, HRV_MedianNN, HRV_MadNN, HRV_MCVNN, HRV_IQRNN, HRV_SDRMSSD, HRV_Prc20NN, HRV_Prc80NN, HRV_pNN50, HRV_pNN20, HRV_MinNN, HRV_MaxNN, HRV_HTI, HRV_TINN, HRV_HF, HRV_VHF, HRV_TP, HRV_HFn, HRV_LnHF, HRV_SD1, HRV_SD2, HRV_SD1SD2, HRV_S, HRV_CSI, HRV_CVI, HRV_CSI_Modified, HRV_PIP, HRV_IALS, HRV_PSS, HRV_PAS, HRV_GI, HRV_SI, HRV_AI, HRV_PI, HRV_C1d, HRV_C1a, HRV_SD1d, HRV_SD1a, HRV_C2d, HRV_C2a, HRV_SD2d, HRV_SD2a, HRV_Cd, HRV_Ca, HRV_SDNNd, HRV_SDNNa, HRV_DFA_alpha1, HRV_MFDFA_alpha1_Width, HRV_MFDFA_alpha1_Peak, HRV_MFDFA_alpha1_Mean, HRV_MFDFA_alpha1_Max, HRV_MFDFA_alpha1_Delta, HRV_MFDFA_alpha1_Asymmetry, HRV_MFDFA_alpha1_Fluctuation, HRV_MFDFA_alpha1_Increment, HRV_ApEn, HRV_ShanEn, HRV_FuzzyEn, HRV_MSEn, HRV_CD, HRV_HFD, HRV_KFD, HRV_LZC, Age, Sex, BMI |
